# Behaviorally emergent hippocampal place maps remain stable during memory recall

**DOI:** 10.1101/2021.07.08.451449

**Authors:** Roland Zemla, Jason J Moore, Jayeeta Basu

## Abstract

The hippocampus is critical for the formation and recall of episodic memories^1, 2^ which store past experience of events (‘what’) occurring at particular locations (‘where’) in time (‘when’). Hippocampal place cells, pyramidal neurons which show location-specific modulation of firing rates during navigation^3, 4^, together form a spatial representation of the environment. It has long been hypothesized that place cells serve as the neural substrate for long-term episodic memory of space^5, 6^. However, recent studies call to question this tenet of the field by demonstrating unexpected levels of representational drift in hippocampal place cells with respect to the duration of episodic memories in mice^7, 8^. In the present study, we examined behaviorally driven long-term organization of the place map, to resolve the relationship between memory and place cells. Leveraging the stability of two-photon calcium imaging, we tracked activity of the same set of CA1 pyramidal neurons during learning and memory recall in an operant, head-fixed, odor-cued spatial navigation task. We found that place cells are rapidly recruited into task-dependent spatial maps, resulting in emergence of orthogonal as well as overlapping representations of space. Further, task-selective place cells used a diverse set of remapping strategies to represent changing task demands that accompany learning. We found behavioral performance dependent divergence of spatial maps between trial types occurs during learning. Finally, imaging during remote recall spanning up to 30 days revealed increased stabilization of learnt place cell maps following memory consolidation. Our findings suggest that a subset of place cells is recruited by rule based spatial learning, actively reconfigured to represent task-relevant spatial relationships, and stabilized following successful learning and consolidation.

## Introduction

The hippocampus plays a critical role in episodic memory formation and recall^9-11^. Across mammalian species, hippocampal place cells show highly specific firing activity for distinct locations in space known as ‘place fields’^3, 4^. While originally hypothesized to serve as a purely navigational mechanism in animals^3^, place cells have since been implicated in displaying mnemonic activity with regard to environmental context^12-14^, object/stimulus association^15, 16^, and trajectory planning^17-19^. Place maps may provide a spatial index (‘where’) for behaviorally-relevant events (‘what’), in service of episodic memory recall.

A central dogma for the cognitive map of space is that, once formed within the structure of episodic memory, hippocampal place maps should retain stable spatial activity previously associated with learning during memory retrieval. *In vivo* electrophysiological recordings in dorsal CA1 in mice performing behaviors with increasing attentional demand to spatial context revealed the greatest increase in the stability of place cell units across 6-hour intervals in mice engaged in cue-dependent navigation^5^. However, long-term imaging studies of place cell activity beyond this interval demonstrated time-limited place cell reactivation^7, 20^ and place field instability^8^ in CA1 when mice were repeatedly exposed to familiar environments. Of note, however, is that none of these studies, however, examined the activity of place cells during operant behaviors that require associational learning with long-term memory demands. Thus, while they clearly show a temporal influence on place map stability, they offer limited understanding of such dynamics under conditions of memory-dependent behavior.

While the hippocampal representations of space must be stable, to enable memory consolidation and recall, they also need to be flexible to allow for accurate discrimination and adaptation of learnt behaviors when environmental contexts change. A key property of place cells is their ability to change their firing fields and/or rates in response to changes in the spatial environment or salient cues within it. This phenomenon is referred to as ‘remapping’^13, 21^. Interestingly, place cells in dorsal CA1 can also remap as a result of learning within a constant spatial environment where task goals, such as reward locations, change but the environment does not. When rats are trained to learn the location of randomly selected reward locations in an open arena, an overrepresentation of place fields emerges in CA1 around the reward areas and the extent of remapping correlates with the performance of the animal^22^. Likewise, goal-oriented learning along a linear treadmill in head-fixed mice induces place cell remapping^23^. Reorganization of spatially tuned ensembles in CA1 occurs during learning of hippocampal-dependent tasks^24^ and is associated with storage of spatial information on both short and long timescales^25^. It is unclear, however, how such spatial maps evolve into orthogonal representations in a context-dependent manner as a result of learnt behavior.

To fill this gap, we used longitudinal two photon imaging in hippocampal CA1 pyramidal neurons while mice learnt and performed a spatial navigation task where they had to collect rewards at distinct specific locations based on different cued contexts. This allowed us to examine mnemonic association of place maps with episodic and contextual task features across learning and recall. Although existing intrinsic hippocampal synaptic mechanisms can lead to the progressive turnover of spatial representations in CA1^7, 22^, we hypothesized that place maps anchored by behavioral learning rules and spatio-temporally structured attentional demands would result in the stabilization and maintenance of learnt task-dependent maps following memory consolidation.

## Results

### Mice learn to reliably perform an odor-cued head-fixed spatial navigation task

To study the time evolution and stability of place cells during episodic memory, we developed a head-fixed, spatial navigation task guided by odor cues. Mice were required to navigate to two discrete unmarked reward zones (10 cm each and separated by ∼80 cm) on a ∼2-meter linear treadmill, each associated with a distinct odor that was delivered at the start of the lap (Extended Data Fig. 1). When the animal received odor A (pentyl acetate), mice had to actively lick in far reward zone A (∼140 cm from lap start, 120 cm from end of odor zone) to receive sucrose water rewards (Fig. 1a). During randomly alternating B trials, when odor B ((+)-α-Pinene) was delivered, rewards became available in the B reward zone (∼60 cm from lap start, 40 cm from end of odor zone) and likewise the animal was operantly rewarded upon licking in this zone (Fig. 1a). Correct learning resulted in the animal licking within the trial-appropriate odor-cued reward zone, while suppressing licking in the alternate trial reward zone. Thus, during an odor A trial, the animal should navigate to and actively lick in reward zone A to get sucrose water rewards, while running past and suppressing licking in B reward zone and its associated anticipatory zone (10 cm prior to reward zone) as well as the rest of the belt; and vice versa for odor B trials. Successful performance thus required the animal to execute a spatial trajectory that depended on memory of the odor cue and sustained spatial attention to execute trial-appropriate lick and lick suppression behavior. Tracking of lick behavior provided us with a reliable readout of learning and subsequent accuracy of episodic memory recall.

**Fig. 1:**
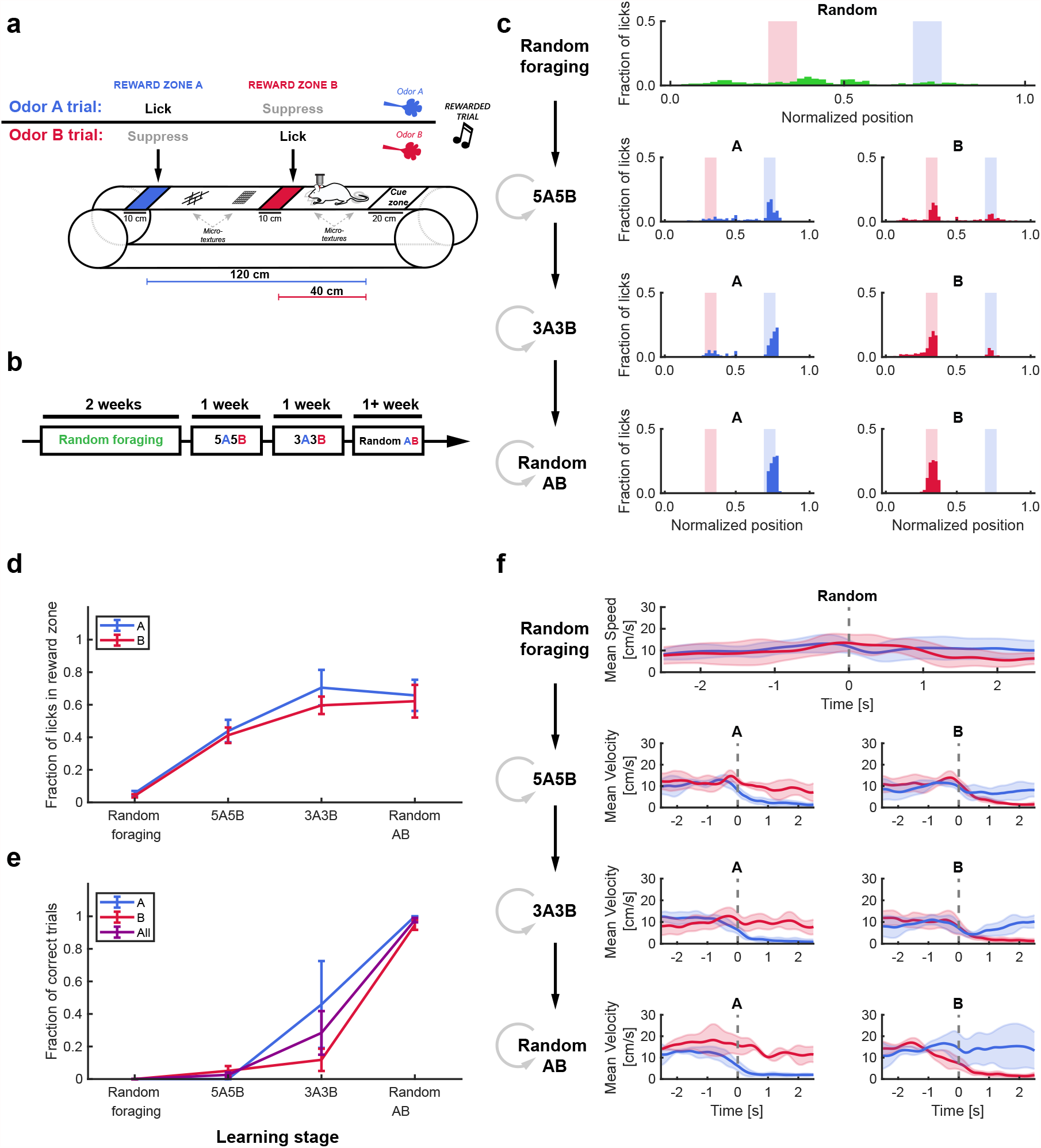
Stable learning of a head-fixed, odor-cued spatial navigation task. (a) Schematic illustrating task structure. During A trials (top row), mice were presented with odor A in a 20 cm cue zone at the onset of the track and ran 120 cm toward the A reward zone (blue patch) to collect 5% sucrose water, while suppressing licking immediately prior to (anticipatory zone) and inside of the B reward zone (red patch) along the way. In contrast, during B trials (bottom row), mice were presented with odor B in the cue zone and ran 40 cm to collect reward within the B reward zone. Following reward collection in B zone, mice were required to suppress lick in the A anticipatory and reward zone for the trial to be registered as correct. Four distinctive micro-textures were placed along the track to aid the mouse in spatial navigation and promote place cell formation. A 2 kHz tone was played immediately prior to the start of each A or B lap to signal that the upcoming trial will be rewarded (as opposed to a non-rewarded time-out lap). (b) Schematic of the timeline for each respective stage of training. Animals were initially trained for 2 weeks to randomly forage, after which they spent ∼1 week on each subsequent stage. (c) Example licking distribution from an animal at each of the four stages of learning. At the random foraging stage, when the animal learned to lick and run, the licking was distributed along the track with no specific enrichment in either the A (blue shade) or B reward zone (red shade). As the animal progressed through each stage of training, licking become more specific to the reward zone associated with that trial (left: A trial laps, right: B trial laps). Circular gray arrows at each stage denote repeated training sessions on different days. (d) The fraction of licks in associated reward zones increased in A and B trials during learning (**fraction of licks, RF vs. Random AB**, paired t-test**: A zone**, 0.06 ± 0.01 vs. 0.66 ± 0.1, \**P* = 0.031; **B zone** 0.04 ± 0.01 vs. 0.62 ± 0.1, \**P* = 0.037; **effect of training stage**, One-way RM ANOVA, **A trials** *F*_1.506, 4.51_ = 12.44, \**P* = 0.017, **B trials** *F*_1.451, 4.354_ = 19.01, \*\**P* = 0.008). (e) Behavioral performance reached >85% only in last stage of training (**fraction of correct trials, RF vs. Random AB**, paired t-test, **A trials**: 0 ± 0 vs. 1 ± 0, \**P* = 0.04, **B trials**, 0 vs. 0.95 ± 0.03, \*\*\**P* < 0.001; **effect of training stage**, one-way RM ANOVA, **A trials**, *F*_1, 3_ = 12.71, *t*_3_ = -∞, p*** < 0.001, **B trials**: *F*_1.004, 3.012_ = 93.36, \*\**P* = 0.002). (f) Example speed of an animal within ± 2 s of entering the reward zones across training stages. At the final training stage (random AB), the animal stops upon entry into a trial-associated reward zone, while running through the non-rewarded zone. Dotted gray line indicates onset time of reward zone entry. Error bars and error shades indicate mean ± s.e.m. Data shown from n = 4 mice.

We implemented a training regimen in which animals were advanced to alternating blocks of serial presentations of A and B trials (stage 1-5A5B; Stage 2-3A3B; Stage 3-random AB) after learning to randomly forage for rewards (Fig. 1b). During random foraging, the licking behavior of the mice was uniformly distributed (Fig. 1c; green subpanel), but with learning in the following stages the licking became more restricted to the respective trial reward zone (Fig. 1c; red and blue subpanels, Fig 1d, Fraction of licks in respective reward zone, stage 1, 2, 3: A trials: 0.06 ± 0.01, 0.44 ± 0.07, 0.71 ± 0.11, 0.66 ± 0.1, *\*\*P* = 0.002; B trials: 0.04 ± 0.01, 0.41 ± 0.05, 0.6 ± 0.05, 0.62 ± 0.1, *\*\*P* = 0.008, mean ± s.e.m, one-way RM ANOVA). Once mice learnt the task, they exhibited consistent trial-appropriate lick and lick suppression behavior indicating acquisition of distinct episodic and context dependent-memories (Fig. 1e). We observed that at this final learnt stage the mice achieve up to 95% accuracy (Fraction of correct trials, stage 1, 2, 3: A trials 0 ± 0, 0 ± 0, 0.46 ± 0.27, 1 ± 0, *\*P* = 0.038; B trials: 0 ± 0, 0.05 ± 0.03, 0.12 ± 0.07, 0.95 ± 0.03, *\*\*P* = 0.002, mean ± s.e.m., one-way RM ANOVA).

To ensure that the animals were navigating toward the trial-appropriate reward zone rather than relying on a dead-reckoning strategy or a texture cue to obtain sucrose water rewards, we measured each animal’s speed immediately prior to and after entry into the reward zones (Fig. 1f, Extended Data Fig. 2). As expected, the animal’s speed decreased immediately prior to (∼1 s) trial-appropriate reward zone entry following learning, but did not decrease in the trial-inappropriate reward zone. Furthermore, animals came to near complete stops within 2 seconds of entering the correct reward zone, suggesting that the animals developed an odor cue-dependent navigational strategy to specific areas of the track. In summary, we developed a head-fixed linear-track based navigation paradigm that allows us to examine episodic, spatial, and contextual features of memory, similar to those previously described in freely moving rodents^17, 19^, which enabled us to study the neural transformation of hippocampal spatial maps under repeated attention and memory demand.

### Task-selective spatial activity emerges in CA1 following learning

Previous studies examining the activity of CA1 pyramidal neurons in animals performing episodic navigation revealed that place firing properties are modulated by past and future behavior of the animal (termed retrospective and prospective coding, respectively)^17-19^. Such neurons (place cells) that show trial-selective activity appear in a space segment common to both trials *prior* to the animal navigating to either the left or right arm of the maze and are hypothesized to facilitate episodic memory recall in the hippocampus. We reasoned that if our task recruits an episodic memory process, we would observe such trial-selective activity along the track segment prior to the mouse licking in either reward zone (i.e. the equivalent common space segment) and possibly beyond.

With this task at hand, we combined the behavior with two photon imaging of Ca^2+^ activity in the cell bodies of CA1 pyramidal neurons endogenously expressing the fluorescent Ca^2+^ indicator GCaMP6f in mice who reached >85% learning criteria (in a cohort of n =10 animals, Fig. 2a). Alongside neurons that were spatially tuned irrespectively of trial type, we observed location specific calcium transients in a sub-population of neurons that were spatially-tuned in a trial-selective manner (Fig. 2b). This means that there was a sub-group of neurons that were place cells on A trials but not on B trials, and vice versa. Examination of the mean run-epoch activity rate (area under curve of calcium transients [AUC]/time) on A vs. B trials revealed that all animals had three groups of neurons which showed preferential activity for either trial (A-selective or B-selective) or shared (A&B) activity (Fig. 2c). We observed that the magnitude of the activity rate was increased more than twofold between laps for preferential A trial and B trial neurons without noticeable difference in activity for non-selective neurons (Fig. 2c, run epochs). Such effect was not observed during epochs when the animal did not run for neither trial-selective neurons nor A&B tuned neurons (Fig. 2c, no-run epochs), suggesting that these neurons were modulated during navigation and incorporated information about the task contingency (Fig. 2c).

**Fig. 2:**
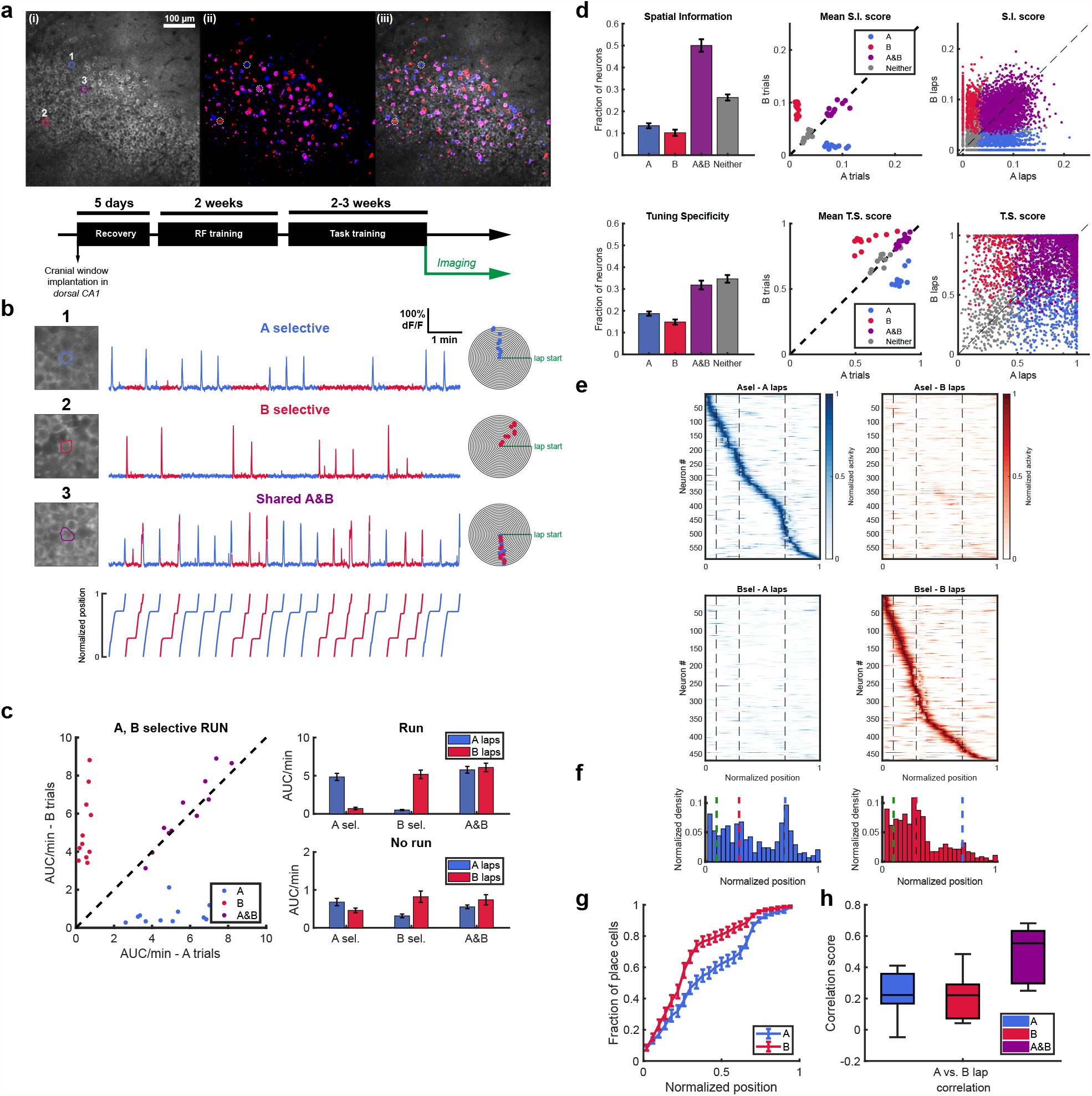
Task-selective place cells are observed in dorsal CA1 during stable performance. (a-i) (Top) Example field of view (FOV) of the pyramidal cell layer imaged in CA1. Image depicts the mean intensity projection. (a-ii) Maximum intensity projection of temporally downsampled run-epoch imaging stacks. Blue overlay represents odor A trials, while red overlay represents odor B trials. Neurons active in both trial types are shown in magenta. (a-iii) Overlay of maximum intensity projection (a-ii) with imaging FOV (a-i). (Bottom) Training timeline of mice. Mice were allowed to recover for 5 days from cranial window surgery and then were trained to run and collect rewards from randomly distributed zones across linear belt (RF – random foraging). Following ∼2 weeks of RF training, mice were transitioned to training on the odor-cued spatial navigation task and imaging began once animals performed at >85% performance for 3 consecutive days. (b) Examples of task selective and non-selective place cells. (1,2) Example of a task-selective place cell on A and B trials, respectively. (3) Example of a non-selective place cell (with place fields on both A and B trials). Numbers correspond to circled neurons in (a). (**c**) (left) Calcium activity rate (AUC [area under curve]/min) of A-, B-, and task non-selective place cells during run epochs. Each point represents the mean from all neurons for each of 11 FOVs from n = 10 animals. (right) Activity of task-selective neurons is greater on their respective task laps during run epochs (**A vs. B lap activity rate, A-selective**: 4.84 ± 0.47 vs. 0.7 ± 0.16, paired Wilcoxon signed-rank test, *W*_10_ = 66, \*\**P* = 0.003; **B-selective**: 0.51 ± 0.07 vs. 5.18 ± 0.55, paired Wilcoxon signed-rank test, *W*_10_ = -66, \*\**P* = 0.003; **non-selective**: 5.77 ± 0.44 vs. 6.08 ± 0.56, paired Wilcoxon signed-rank test, *W*_10_ = -30, *P* = 0.206), while no difference is observed during no run epochs (**A-selective**: 0.68 ± 0.1 vs. 0.46 ± 0.06, paired Wilcoxon signed-rank test, *W*_10_ = 46, *P* = 0.082; **B-selective** 0.31 ± 0.05 vs. 0.82 ± 0.15, paired Wilcoxon signed-rank test, *W*_10_ = -62, \*\**P* = 0.009; **non-selective**: 0.55 ± 0.05 vs. 0.74 ± 0.13, paired Wilcoxon signed-rank test, *W*_10_ = -32, *P* = 0.175). (d) (left) Fraction of place cells tuned according to spatial information (S.I.) and tuning specificity (T.S.) showed distinct distributions between trials (**S**.**I. fraction**: Friedman test, 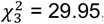, \*\*\**P* < 0.001; **T**.**S. fraction**: Friedman test, 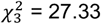, \*\*\**P* < 0.001). More A-than B-selective neurons were generally present using the S.I. score (**fraction of A vs. B**: 0.13 ± 0.01 vs. 0.1 ± 0.01, paired Wilcoxon signed-rank test, *W*_10_ = 54, \**P* = 0.014) and T.S. score (0.19 ± 0.01 vs. 0.15 ± 0.01, paired Wilcoxon signed-rank test, *W*_10_ = 48, \**P* = 0.032). Both A- and B-selective were fewer in number compared to A&B neurons using the S.I. score (**A vs. A&B**: 0.13 ± 0.01 vs. 0.5 ± 0.03, paired Wilcoxon signed-rank test, *W*_10_ = -66, \*\**P* = 0.003; **B vs. A&B**: 0.1 ± 0.01 vs. 0.5 ± 0.03, paired Wilcoxon signed-rank test, *W*_10_ = - 66, \*\**P* = 0.003) and T.S. score (**A vs. A&B**: 0.19 ± 0.01 vs. 0.32 ± 0.02, paired Wilcoxon signed-rank test, *W*_10_ = -66, \*\**P* = 0.003; **B vs. A&B**: 0.15 ± 0.01 vs. 0.32 ± 0.02, paired Wilcoxon signed-rank test, *W*_10_ = -66, \*\**P* = 0.003). Error bars represent mean ± s.e.m. (center). Mean spatial information scores (bits/Ca^2+^ event) and tuning specificity score for each class of neurons on A and B laps during each session. (right) Spatial information and tuning specificity scores from all imaged neurons (5158 neurons). (e) Rate maps from all the mice for A- and B-task selective place cells on A and B laps. The rate of each neuron is normalized to its maximum rate across both trial types. A-selective neurons (top) are sorted according to their maximum rate across 100 spatial bins on A laps. The same sorting was performed on B-selective neurons (bottom) on B laps. Green dashed line indicates the end of odor zone, red the start of B reward zone, and blue the start of A reward zone. (f) Distribution of the place field centroid for A-selective and B-selective place cells across the track (25 spatial bins). Both categories of place cells are non-uniformly distributed across the track, with a skew toward the common segment of the track (**place field distribution, A-selective**: Rayleigh test of uniformity, *Z* = 4.72, \*\**P* = 0.009, *n* = 590 neurons; **B-selective**: Rayleigh test of uniformity, *Z* = 89.63, \*\*\**P* < 0.001, n = 468 neurons). A-selective place cells tend to be also distributed at toward more distant locations on the track toward the A reward zone. (**g**) Distribution of place field centroids differs between A-selective and B-selective place cells (**A vs. B place field centroid difference**: 2-sample Kolmogorov-Smirnov test, *D*_590, 468_ = 0.22, \*\*\**P* < 0.001, *n* = 590 vs. 468 neurons). (**h**) Pearson correlation of spatial tuning curves between A and B laps for A-, B-, and trial non-selective place cells. Spatial correction scores are low for task selective neurons with no difference between groups and significantly lower compared to task non-selective neurons consistent with effective discrimination between each category of place cells (**A-selective vs. A&B**: 0.24 ± 0.04 vs. 0.49 ± 0.05, paired Wilcoxon signed-rank test, *W*_10_ = -66, \*\**P* = 0.003; **B-selective vs. A&B**: 0.21 ± 0.04 vs. 0.49 ± 0.05, paired Wilcoxon signed-rank test, *W*_10_ = -66, \*\**P* = 0.003). Central mark indicates median and top and bottom boxes indicate 25^th^ and 75^th^ percentiles, respectively. Whiskers denote the most extreme data points. Error bars indicate mean ± s.e.m. Data shown from 11 FOV from n = 10 mice.

To examine the task-dependent spatial tuning of neurons, we used two previously described metrics to detect place cells: spatial information (S.I.) and tuning specificity (T.S.)^23, 26^. Each metric has its unique advantage and we used both to maximize detection accuracy. While spatial information allows for detection of place cells with single- and multiple place field, it has lower sensitivity for cells with broad, single fields. On the other hand, tuning specificity is more sensitive (and more specific) to place cells with single fields regardless of field width at the expense of multi-field cell detection^23^. Regardless of the place cell classification criteria, each metric revealed the existence of A-, B-selective as well as trial-nonselective (A&B) place cells in similar fractions as expected from the activity rate analysis above (Fig. 2d). We noted more A-selective than B-selective place cells using the S.I. metric and observed a similar trend using the T.S metric (Fig. 2d), perhaps due to the greater length of the track traversed to arrive at the A-reward goal location. Using either spatial tuning metric, there were nearly half as many B-selective neurons as A&B neurons and we similarly observed this trend for A-selective neurons compared to A&B neurons (Fig. 2d). We combined both tuning criteria to determine the task-selectivity of place cells (i.e. a selective place cell required tuning by either criterion in one set of trials and not in the alternate trials). We did not observe significant speed differences between trials in all the animals we recorded from, with a mean difference in speed not exceeding ± 5 cm/s in all spatial bins except for peri-reward bins (Extended Data Fig. 3). In neurons with task-selective place fields, the majority of place cells did not show a significant speed difference suggesting that the selectivity we observed is not due to an effect of speed on Ca^2+^ activity between trials (Extended Data Fig. 4a, b). Furthermore, we could not attribute such difference to the sensitivity of Ca^2+^ transient detection between trials as GCaMP6f detects single action potentials *in vivo*^*27*^ and our detection algorithm is optimized for detection of small transients (see Methods). Taken together, the presence of trial-selective place cells suggests episodic encoding of space in our task.

The task-selective, and thus episodic, nature of place cell activity in our task suggested that the distribution of place fields along the track would also vary between trials due to the distinct memory and attention demands associated with navigation to each reward zone. Both A- and B-selective place cells had place fields that spanned the entire length of the track (Fig. 2e). However, there was a significant difference in the distribution of place fields within A trials and B trials for A-only, and B-only task-selective neurons (Fig. 2f). The difference in place field location distribution was also significant between A- and B-selective neurons (Fig. 2g). We observed an overabundance of place fields within the common track segment – from lap start until the B reward zone – on both trials consistent with the greatest behavioral significance of this area in navigating a trial trajectory (near (B) or far (A) zone destination). Interestingly, there was a greater density of place fields in this segment on B trials with a rapid decline of field density thereafter in agreement with a shorter B trajectory. In contrast, the distribution of the longer trajectory A-selective place fields was more uniform across the track with increasing field density near the distant reward zone. This may imply that the ‘near’ reward zone B is significant in guiding behavioral output choices in both trials, as the mouse either has to stop and lick for rewards at the near B reward zone for Odor B trials, whereas actively suppress its licking to cross over that B zone to seek rewards in the far A zone for Odor A trials. We did not observe a difference in the place field properties between trials (Extended Data Fig. 4c, d). Lastly, we compared the spatial tuning curve correlations between A and B trials for A- and B-selective place cells against shared A&B place cells. We observed low correlation scores for A-selective and B-selective neurons with no significant difference between them (Fig. 2h), indicating that activity maps from A- and B-selective neurons discriminate between trial types. Significantly higher correlation scores were present among the A&B shared place cells relative to A-selective and B-selective neurons (Fig. 2h). Furthermore, we observed a broad range of correlation scores among the task-nonselective group – with some neurons approaching scores similar to those of the task selective neurons – suggesting that more complex trial-to-trial spatial dynamics exist among this subpopulation.

### Place cells exhibit dynamic, behaviorally-driven remapping properties between trial types

The activity of place cells is most prominently modulated by the location of the animal in the environment. However, changes in the sensory environment and behavioral demands can influence the activity of place cells as well^12, 13, 17^-^19, 21, 28, 29^. This property has previously been described as ‘remapping’ and is expressed by changes in place field firing rate and/or place field location, also known as rate and global remapping, respectively^13^. As the sensory environment in our task remained fixed (except for changes in odor identity at the lap start), any remapping activity between A and B trials would reflect behavioral modulation of spatial maps and suggests episodic encoding.

To examine the remapping features of A&B shared place cells, we classified each neuron into one of four remapping categories according to their cross-trial spatial tuning curve similarly and place field-related calcium activity (Fig. 3a, b). Based on commonly used nomenclature in the field established through electrophysiology studies ^13^, we defined three classes of remapping neurons: global, activity (in lieu of rate), and partial remapping (Fig 3b). Global remapping neurons had consistent shifts in place field location between trials identified by the dissimilarity of their A and B tuning curves (Extended Data Fig. 5, 6). Among the population of common field place cells (common), we identified a subset of neurons whose calcium activity was modulated by trial type that we labeled as activity remapping. These neurons were analogous to firing rate remapping neurons described in *in vivo* electrophysiological studies^13^. We used a peak activity modulation index to verify that these neurons indeed represented a subset of common place cells (Extended Data Fig. 7). Lastly, we identified a unique population of neurons that exhibited what we termed partial remapping. These neurons shared a common place field between trials, but had an additional place field unique to one trial type (i.e. partially remapped). All classes of remapping cells were present along the entire length of the track (Fig. 3c). The greatest fraction of place cells was in the non-remapping common category, followed by global and partial remapping neurons with a minority of place cells classified as activity remapping (Fig. 3d). We observed a significant difference between the fraction of common and activity remapping neurons and common and partial remapping neurons, while no difference between common and global remapping neurons (Fig. 3d).

**Fig. 3:**
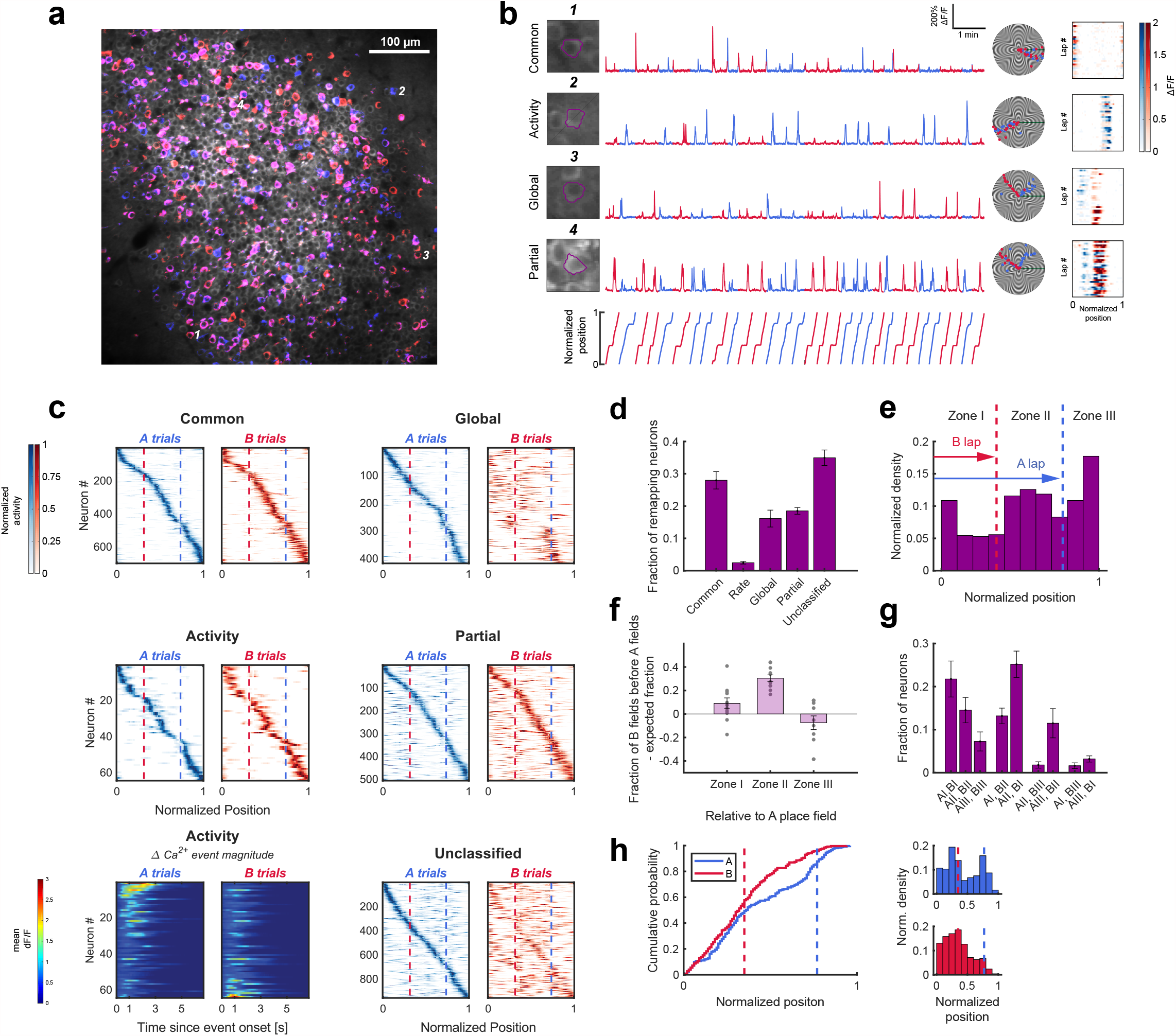
Place cells show distinct and task-oriented forms of remapping between trial types. (a) Overlap of the imaging field with the maximum intensity projection on A-laps (blue) and B-laps (red). (b) Examples of the three types of remapping place cells observed in CA1. (1) Example of a common neuron that fires in its place field regardless of trial type. (2) Example of an activity remapping neuron whose calcium activity in its place field is modulated by trial type. (3) Example of a global remapping neuron which has distinct place fields on each trial type. (4) Examples of a partially remapping neuron with a common field (trial insensitive) located ∼1/3 of the distance from the lap start and an A trial specific field (partial field) located before it. Individual points on event spiral maps represent significant running-related Ca^2+^ events on A (blue) or B (red) trials. Colormaps on the right represent the mean ΔF/F activity in each spatial bin on each of the pseudo-randomly presented trial laps. Blue colormap represents A trials, while red colormap represented B trials. Note the difference in the ΔF/F signal of the activity remapping (2) neuron between A and B laps. Example numbers correspond to the neurons circled in (a). (c) Spatial tuning colormaps for each class of remapping place cells. Cells are sorted according to the maximum spatial bin rate on A laps. Note the predominant shift of global remapping neurons place fields toward earlier locations on the track on B laps. Partial remapping neurons were sorted by their common place field. Bottom left panel depicts the mean ΔF/F value relative to the onset of the Ca^2+^ event in the place field for activity remapping neurons. The activity map was sorted according to the difference between the peak mean ΔF/F value of Ca^2+^ transients in the place field on A vs. B trials to emphasize the degree of rate remapping in contrast to the spatial tuning map above in which place cells are sorted according to their maximum spatial bin rate on A laps. The unclassified category consists of spatially tuned neurons that had 2+ place field on both trials. (d) Distribution of the classes of non-remapping (common), remapping, and unclassified place cells. The difference in distribution was significant between the common class and three classes of remapping neurons (**Difference among remapping classes**: Friedman test, 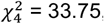, \*\*\**P* < 0.001; **Common vs. activity**: 0.28 ± 0.03 vs. 0.02 ± 0, paired Wilcoxon signed-rank test, *W*_10_ = 66, \*\**P* = 0.003; **Common vs. global**: 0.28 ± 0.03 vs. 0.16 ± 0.03, paired Wilcoxon signed-rank test, *W*_10_ = 48, \**P* = 0.032; **Common vs. partial**: 0.28 ± 0.03 vs. 0.19 ± 0.01, paired Wilcoxon signed-rank test, *W*_10_ = 60, \*\**P* = 0.01). Error bars indicate mean ± s.e.m. (e) Common place cells are distributed across the track according to spatial task demand. The lowest density of common fields was present in Zone I (prior to the B reward zone) where spatial orientation/mapping is critical to task performance. The place field density progressively increased from Zone I until zone III (**Place field distribution for common neurons**: Rayleigh test of uniformity, *Z* = 13.82, \*\*\**P* < 0.001, *n* = 700 neurons). (f) Global remapping neurons exhibited task-oriented remapping of place fields. In particular, place cells with place field centroids located in Zone II (between the reward zones) on A laps exhibited a statistically significant shift of place fields on B trials to earlier positions on the track (**Zone II A vs. B lap field shift:** 0.3 ± 0.03, 1-sample Wilcoxon signed-rank test against 0, *W*_10_ = 66, \*\*\**P* < 0.001). (g) Analysis of inter-zone movement revealed additional shifting of A place fields relative to B fields (**Global remapping neurons place field zone shift**: Friedman test, 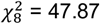, \*\*\**P* < 0.001) There were, however, no significant shifts between specific zones. Error bars indicates mean ± s.e.m. (h) Partial fields of partial remapping place cells exhibited remapping dynamics similar to those observed for task-selective place cells (Figure 2f) with fields skewed toward zone I of the track and additional fields in Zone II during A trials (**A vs. B partial remapping neurons place field centroid difference**: 2-sample Kolmogorov-Smirnov test, *D*_253, 253_ = 0.19, \*\*\**P* < 0.001, *n* = 253 vs. 253 neurons). Arrows represent the respective reward zones. Analysis from 11 FOV from n = 10 mice.

Given our observation of task-dependent place cell remapping, we wanted to ask whether the pattern of spatial remapping correlated with the behavioral demands of each trial type as we observed for task selective place cells. Our task design introduced an implicit behavioral gradient with progressively lower memory and attention demands once the animal had traversed the directed goal location, collected the appropriate reward and ran toward the end of the track. To link place coding with this behavioral gradient in our analysis, we split the track into three spatial zones defined by the reward zones and first examined the distribution of common (non-remapping) place cells. Common place cells showed a decrease in field density with zone distance (Fig. 3e). Similar to the distribution of A- and B-selective place cell fields (Fig. 2f), the distance of the trial trajectories was also conveyed by global remapping place cells with place cells in zone II on longer trajectory A trials shifting their place fields to earlier locations on shorter trajectory B trials (Fig. 3f). In contrast, such remapping neither occurred in Zone I nor Zone III (Fig. 3f). We also observed an inter-zonal, near significant, tendency for global place cells to preferentially shift their place fields from Zone II on A trials to Zone I on B trials rather than in the opposite direction and a significant shift of Zone III A place fields to Zone II on B trials (Fig. 3g). Shifting of place fields within the same zones occurred at the same frequency in all three zones (data not shown). Partial remapping neurons likewise showed a trial-dependent trajectory distribution of trial-specific fields with an overrepresentation of fields in Zone I for both trial types and an additional overrepresentation of fields around the more distant reward zone on A trials (Fig. 3h).

We also observed that partial remapping neurons with common place fields in Zone II had an overrepresentation of B trial-specific at earlier locations on the track (data not shown).

### Task-dependent spatial maps retain stable activity after learning during recall

Next, we asked whether spatial maps emerging during the task remain stable as the memory is acquired and consolidated. Previous one-photon and two-photon imaging experiments of CA1 place cells reported a rapid decorrelation of place maps over several days^8, 23^ under little or variable memory demand. However, *in vivo* electrophysiological recordings in CA1 revealed a strong association between spatial map stability and memory- and attention-dependent navigation at 6 hours^5^. To determine whether place maps are more stable when learnt within the structure of a memory task at longer timescales, we compared the place maps of the same CA1 pyramidal neurons imaged during an accelerated training regimen (learning) against place maps acquired following consolidated learning (recall) (Fig. 4a). Mice learnt the task with 92 ± 2% accuracy (mean ± s.e.m.; n = 6) by day 7 on the accelerated regimen (Fig. 4b), while performance during recall remained consistently between ∼80-100% (Fig. 5d, performance plot). We tracked neurons using an automated ROI registration algorithm as part of the CaImAn analysis package^30^ and manually discarded low quality matches (Fig. 4c, Extended Data Fig. 8; see Methods for details).

**Fig. 4:**
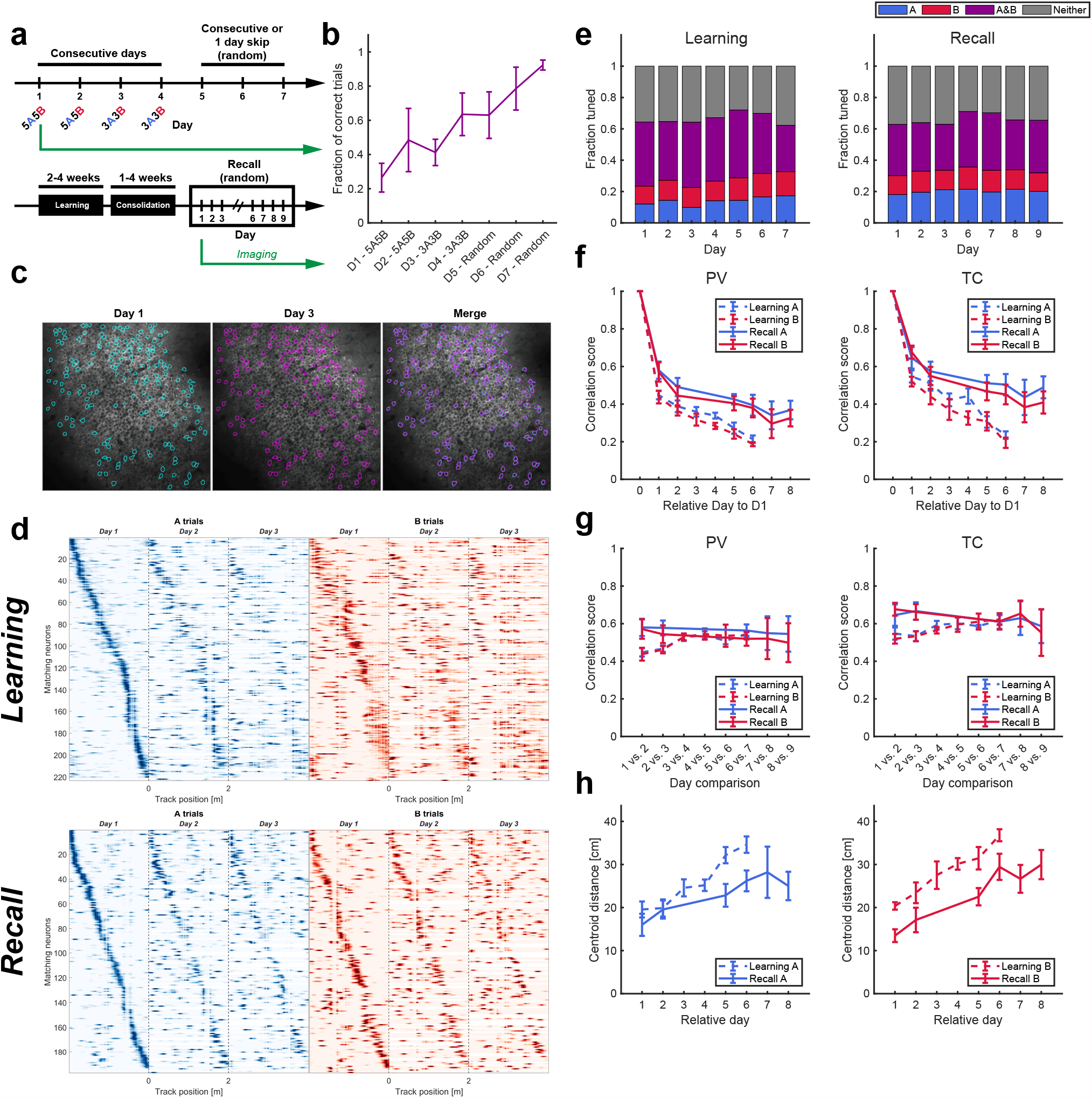
Learning induces rapid remapping of place maps. (a) (Top) Schematic illustrating the accelerated training regimen and imaging schedule for odor-cued spatial navigation. (Bottom) Timeline for recall imaging experiments following memory consolidation. (b) Learning performance during accelerated learning. Mice achieve high behavior performance by the last day of training (92 ± 2%, *n* = 6 mice). Error bars indicate mean ± s.e.m. (c) Example of matching spatial components (ROIs) across two different sessions. Only components with high spatial component overlaps were used in the analysis to avoid component mismatching. (d) (Top) Example spatial tuning maps that show rapid remapping of spatial activity during learning in contrast to memory recall following consolidation (Bottom). (e) Task-dependent tuning of place cells occurs as early as the first day of training and persists during learning with increase in fraction of A-trial tuned place cells during A, but not B trials (**Fraction of tuned place cells, A trials, learning**: one-way RM mixed effects analysis, effect of training day, *F*_6, 22.24_ = 7.24, \*\*\**P* < 0.001; **B trials**: one-way RM mixed effects analysis, effect of training day, *F*_6, 27_ = 1.36, *P* = 0.265, *n* = 6 mice) with no change during recall (**A trials, recall**: one-way RM mixed effects analysis, effect of training day, *F*_6, 28_ = 0.32, *P* = 0.923;**B trials**: one-way RM mixed effects analysis, effect of training day, *F*_6, 28_ = 0.26, *P* = 0.951, *n* = 5 mice) (f) (left) Population vector (PV) correlation of all matching cells relative to Day 1 of imaging shows rapid restructuring of run-related activity on the following training day that stabilizes on recall trials (**PV correlation, A trials**: two-way RM mixed effects analysis, effect of time, *F*_3, 22.35_ = 50.93, \*\*\**P* < 0.001, effect of behavior, *F*_1, 9.04_ = 9.02, \**P* = 0.015, interaction between time and behavior, *F*_3, 22.35_ = 0.78, *P* = 0.519; **B trials**: two-way RM mixed effects analysis, effect of time, *F*_3, 22.44_ = 43.88, \*\*\**P* < 0.001, effect of behavior, *F*_1, 9.07_ = 9.13, \**P* = 0.014, interaction between time and behavior, *F*_3, 22.44_ = 1.78, *P* = 0.18, *n* = 6 learn, 5 recall mice; **Day 2 vs. 7, recall, A trials**: 0.58 ± 0.05 vs. 0.39 ± 0.06, paired *t*-test, *t*_4_ = 7.89, \*\**P* = 0.003; **B trials**: 0.57 ± 0.05 vs. 0.38 ± 0.05, paired *t*-test, *t*_4_ = 7.79, \*\**P* = 0.002, *n* = 5 mice), but not during learning (**Day 2 vs. 7, learning, A trials**: 0.43 ± 0.03 vs. 0.19 ± 0.01, paired *t*-test, *t*_2_ = 5.89, \**P* = 0.028; **B trials**: 0.41 ± 0.04 vs. 0.19 ± 0.01, paired *t*-test, *t*_2_ = 5.06, *P* = 0.053, *n* = 3 mice; **Day 7, learning vs. recall, A trials**: 0.21 ± 0.02 vs. 0.39 ± 0.06, unpaired *t*-test, *t*_7_ = -2.78, \**P* = 0.027; **B trials**: 0.19 ± 0.01 vs. 0.38 ± 0.05, unpaired *t*-test, *t*_7_ = -3.37, \**P* = 0.014, *n* = 4 learn, 5 recall mice). A similar trend was observed for the **tuning curve** correlation scores between matching place cells selected using the tuning specificity criterion (**TC correlation, A trials**: two-way RM mixed effects analysis, effect of time, *F*_3, 22.37_ = 27.73, \*\*\**P* < 0.001, effect of behavior, *F*_1, 8.85_ = 8.69, \**P* = 0.017, interaction between time and behavior, *F*_3, 22.37_ = 3.95, \**P* = 0.021; **B trials**: two-way RM mixed effects analysis, effect of time, *F*_3, 22.46_ = 32.87, \*\*\**P* < 0.001, effect of behavior, *F*_1, 8.81_ = 11.81, \*\**P* = 0.008, interaction between time and behavior, *F*_3, 22.46_ = 1.73, *P* = 0.189, *n* = 6 learn, 5 recall mice; **Day 2 vs. 7, learning, A trials**: 0.51 ± 0.03 vs. 0.22 ± 0.02, paired *t*-test, *t*_2_ = 5.84, \**P* = 0.028; **B trials**: 0.51 ± 0.03 vs. 0.19 ± 0.04, paired *t*-test, *t*_2_ = 6.49, \**P* = 0.045, *n* = 3 mice; **Day 2 vs. 7 recall, A trials**: 0.65 ± 0.06 vs. 0.5 ± 0.06, paired *t*-test, *t*_4_ = 5.1, \**P* = 0.014; **B trials**: 0.68 ± 0.03 vs. 0.45 ± 0.05, paired *t*-test, *t*_4_ = 5.32, \*\**P* = 0.006, *n* = 5 mice; **Day 7, learning vs. recall, A trials**: 0.23 ± 0.02 vs. 0.5 ± 0.06, unpaired *t*-test, *t*_7_ = -4.05, \*\**P* = 0.009; **B trials**: 0.19 ± 0.03 vs. 0.45 ± 0.05, unpaired *t*-test, *t*_7_ = -4.03, \*\**P* = 0.01, *n* = 4 learn, 5 recall mice). (g) Learning stabilizes neighboring session maps at the population level and between place cells. As learning progresses through each training stage, the population correlation scores approach those observed for the memory-consolidated recall cohort (**Neighboring session PV correlation, A trials**: two-way RM mixed effects analysis, effect of time, *F*_2, 14.28_ = 3.98, \**P* = 0.042, effect of behavior, *F*_1, 9.24_ = 5.45, \**P* = 0.044, interaction between time and behavior, *F*_2, 14.28_ = 7.37, \*\**P* = 0.006; **B trials**: two-way RM mixed effects analysis, effect of time, *F*_2, 14.17_ = 0.98, *P* = 0.4, effect of behavior, *F*_1, 9.06_ = 1.88, *P* = 0.204, interaction between time and behavior, *F*_2, 14.17_ = 6.18, \**P* = 0.012, *n* = 6 learn, 5 recall mice; **Days 1 vs. 2 Vs. Day 6 vs. 7, learning, A trials**: 0.43 ± 0.03 vs. 0.54 ± 0.01, paired *t*-test, *t*_2_ = -5.24, \**P* = 0.034; **B trials**: 0.41 ± 0.04 vs. 0.52 ± 0.01, paired *t*-test, *t*_2_ = -2.42, *P* = 0.136, *n* = 3 mice; **Days 1 vs. 2 Vs. Day 6 vs. 7, recall, A trials**: 0.58 ± 0.05 vs. 0.56 ± 0.03, paired *t*-test, *t*_4_ = 0.55, *P* = 0.611; **B trials**: 0.57 ± 0.05 vs. 0.52 ± 0.04, paired *t*-test, *t*_4_ = 1.54, *P* = 0.198, *n* = 5 mice). For spatially tuned neurons, we the stabilization occurred during learning on both A and B trials, but not during recall (**Neighboring session TC correlation, A trials**: two-way RM mixed effects analysis, effect of time, *F*_2, 14.38_ = 0.74, *P* = 0.492, effect of behavior, *F*_1, 9.14_ = 2.32, *P* = 0.161, **interaction between time and behavior**, *F*_2, 14.38_ = 6.01, \**P* = 0.013; **B trials**: two-way RM mixed effects analysis, effect of time, *F*_2, 13.98_ = 0.55, *P* = 0.586, effect of behavior, *F*_1, 8.83_ = 4.49, *P* = 0.064, **interaction between time and behavior**, *F*_2, 13.98_ = 9.32, \*\**P* = 0.003, *n* = 6 learn, 5 recall mice; **Days 1 vs. 2 Vs. Day 6 vs. 7, learning, A trials**: 0.51 ± 0.03 vs. 0.61 ± 0.03, paired *t*-test, *t*_2_ = -5.89, \**P* = 0.028; **B trials**: 0.51 ± 0.03 vs. 0.6 ± 0.01, paired *t*-test, *t*_2_ = -5.09, \**P* = 0.036, *n* = 3 mice; **Days 1 vs. 2 Vs. Day 6 vs. 7, recall, A trials**: 0.65 ± 0.06 vs. 0.61 ± 0.04. paired *t*-test, *t*_4_ = 0.81, *P* = 0.465; **B trials**: 0.68 ± 0.03 vs. 0.61 ± 0.04, paired *t*-test, *t*_4_ = 1.71, *P* = 0.162, *n* = 5 mice). (h) Place fields are remapped over greater distances during learning on A and B trials (**Mean centroid difference relative to Day 1, A trials**: two-way RM mixed effects analysis, effect of time, *F*_3, 22.19_ = 30.7, \*\*\**P* < 0.001, effect of behavior, *F*_1, 8.62_ = 3.52, *P* = 0.095, interaction between time and behavior, *F*_3, 22.19_ = 2.78, *P* = 0.065, *n* = 6 learn, 5 recall mice; **B trials**: two-way RM mixed effects analysis, effect of time, *F*_3, 23.34_ = 25.56, \*\*\**P* < 0.001, effect of behavior, *F*_1, 8.98_ = 11.81, \*\**P* = 0.007, interaction between time and behavior, *F*_3, 23.34_ = 0.14, *P* = 0.936, *n* = 6 learn, 5 recall mice; **Learning vs. recall, Day 5, A trials**: 32.16 ± 1.89 cm vs. 22.83 ± 2.64 cm, unpaired *t*-test, *t*_8_ = 2.88, \**P* = 0.041; **B trials**: 31.45 ± 2.6 cm vs. 22.51 ± 2.01 cm, unpaired *t*-test, *t*_8_ = 2.72, *P* = 0.052, *n* = 5 learn, 5 recall mice).

Analysis of spatial activity of neurons revealed instability of the network during task learning in the familiar spatial environment, while the place map network showed remarkable stability during the recall sessions (Fig. 4d). To quantify spatial task selectivity across days, we selected all cells that were significantly tuned according to the tuning specificity (T.S.) criterion on a given session and compared their distribution across time. We chose the T.S criterion to favor the selection of place cells with single place fields. Using place cells classified using the spatial information (S.I.) metric yielded similar results (Extended Data Fig. 9). As early as the first day of training, we observed trial-selective tuning on A and B trials and a subtle, but statistically insignificant increase in the fraction of task-selective neurons across time at 6 and 7 days from the start of imaging on A and B trials (Fig. 4e). During recall stage, the task-specific distribution of place cells remained stable across time on A and B trials (Fig. 4e). Importantly, we observed a significant difference in the stability of place maps at both the population level and individual place cells matched to the first day of imaging. The population vector (PV) correlation showed a decorrelation for both learning A and B trials as well as recall A and B trials, but was substantially lower for the learning cohort on day 6 and day 7 compared to recall for A and B trials (Fig. 4f). When we specifically looked at place cells, we observed a rapid tuning curve decorrelation beginning at day 2 for the learning cohort that continued over time on A and B trials as well as for the recall cohort on A and B trials (Fig. 4f). However, the correlation scores on day 6 and 7 on A and B trials were significantly lower during learning compared to recall suggesting a stabilization of spatial representations following learnt consolidation of memory in the recall cohort.

To examine the rate at which the spatial network stabilized, we calculated the correlation between neighboring day sessions at the level of the population and individual place cells (Fig. 4g). We reasoned that as the animals learnt the task rules the activity maps would reach neighboring day scores similar to that of the recall cohort, which we expected to be fixed following consolidation. We observed that at the population level (PV score), the learning cohort experienced a time-dependent increase in neighboring day map similarity, but this was not present in the recall cohort on A or B trials (Fig. 4g left). In contrast, spatial maps in the learning cohort increased their neighboring day correlation (TC score) across time on *both* A and B trials, whereas we did not observe such increase in the recall animals on either A or B trials (Fig. 4g, right).

To determine whether there is a shift in the place field locations, as a correlate for reconfiguration of the network in each trial type, we calculated the mean change of the place field centroids relative to the first imaging session. For A trials, we did not observe a mean change of centroid location relative to recall animals until day 3 and observed a progressive increase in the distance of place field centroid remapping thereafter (Fig. 4h). In contrast, B trial place cells maintained a consistently higher rate of place field centroid shift throughout learning (Fig. 4h). Lastly, we imaged a subset of recall animals (n = 3) over longer term to examine how long can the activity of learnt spatial maps persist. We observed a surprisingly high level of map stability across 30 days of imaging, with a progressive, albeit slow, decay in correlation (Extended Data Fig. 10).

Overall, our imaging findings support the idea that learning induces place remapping that stabilizes in the long-term during recall phases when memory is consolidated.

### Dissimilarity between task-specific spatial maps predicts performance

While task-modulated place cells appear to be a feature of episodic spatial behavior, the link between learning and the activity of these maps remains unclear. Inactivation experiments have shown that the activity of these cells is not necessary for successful performance on a continuous T-arm alternation maze^31^. On the other hand, shorter-timescale (during recall) inactivation of CA1 pyramidal neurons has causally linked place cell maps to memory consolidation ^32^. To understand how strongly the rate of learning is coupled to the rate of task-dependent spatial map divergence, we compared the A vs. B spatial tuning correlation of place cells tuned in both trial laps (i.e., A&B tuned) during learning and recall relative to the first day of imaging across time. We noticed a greater tendency for place cells to split their place fields between tasks during learning (Fig. 5a) than during recall (Fig. 5b). When we quantified this effect, we observed a significant decorrelation of the tuning curves of A&B tuned place cells which progressively increased (i.e. correlation decreased) as the performance of the animals increased during learning (Fig. 5c). This effect was observed as early as the second day of imaging and continued until day seven (Fig. 5c). Interestingly, we did not see a similar decorrelation trend between A&B place cells during recall experiments (Fig. 5d). Furthermore, we quantified the A vs. B correlation in all place cells as a function of animal performance across all the imaged sessions and observed a strong inverse correlation during learning (Fig. 5e). In contrast, we did not observe a significant correlation during recall (Fig. 5f). Thus, existing spatial maps are remapped to reflect the degree of task learning in CA1.

**Fig. 5:**
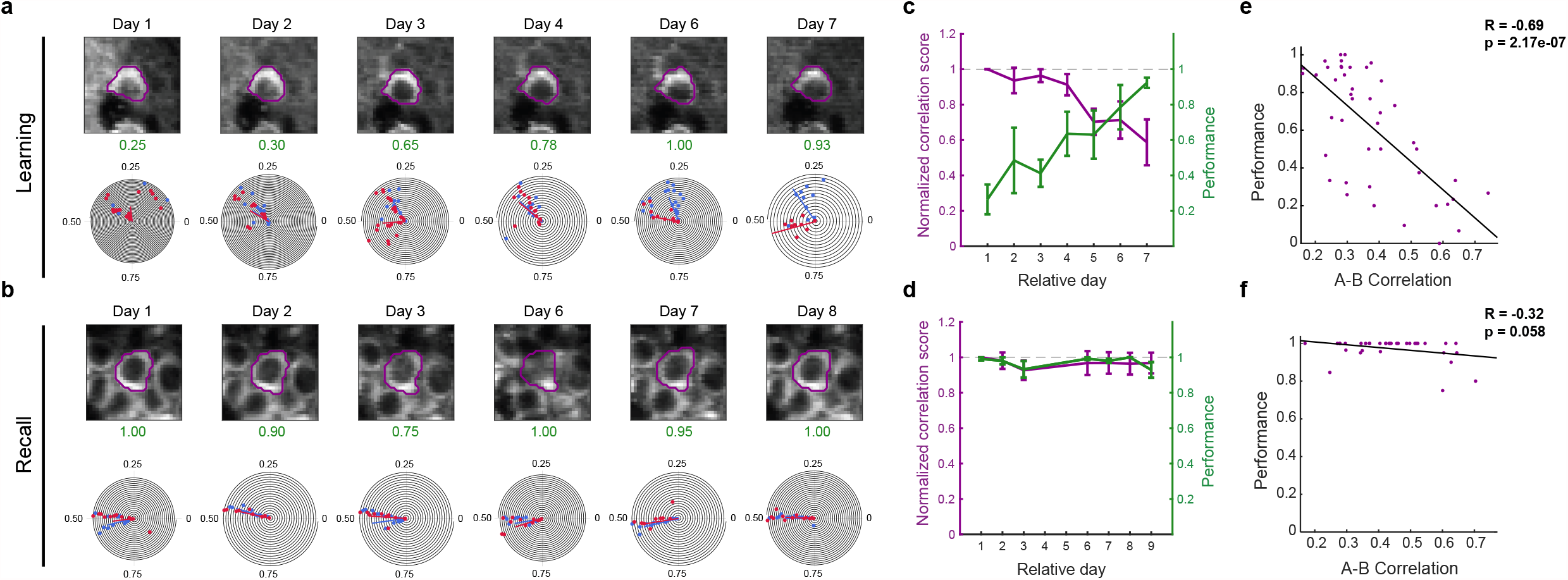
Dissimilarity between task spatial maps predicts animal performance. (a) Example neuron during learning. As the performance of the animal increases during training sessions (green), the task non-selective neuron ‘splits’ its common place field into two task-dependent fields. (b) During recall, the spatial fields of this neurons are stable. (c-d) Overlay of plots of normalized tuning curve correlation scores (left axis) and animal performance (right axis) of matched A&B spatially tuned neurons during learning (c) and recall (d). Normalized tuning curve correlation scores were calculated as the A lap vs. B lap tuning correlation score on the relative session day divided by the A vs B lap correlation score on the first day of imaging for each matched neuron. As the performance of animals increased during learning, place cell maps between task trials became progressively more decorrelated (**Normalized A vs. B lap correlation scores, learning**: Kruskal-Wallis test, *H*_5_ = 63.31, \*\*\**P* < 0.001, *n* = 1050 neurons from 6 mice; **Day 2**: 0.94 ± 0.07, 1-sample Wilcoxon signed-rank test against 1, *W*_178_ = -3536, \**P* = 0.011, *n* = 179 neurons; **Day 7**: 0.59 ± 0.13, 1-sample Wilcoxon signed-rank test against 1, *W*_121_ = -4197, \*\*\**P* < 0.001, *n* = 122 neurons). A similar effect was not observed during recall when animals performed >90% accuracy on each day and the normalized correlation scores were near the expected value of 1 (dashed gray line; **Normalized A vs. B lap correlation scores, recall**: Kruskal-Wallis test, *H*_5_ = 7.15, *P* = 0.21, *n* = 577 neurons from 5 mice; **Day 2**: 0.99 ± 0.05, 1-sample Wilcoxon signed-rank test against 1, *W*_132_ = -5, *P* = 0.996, *n* = 133 neurons; **Day 7**: 0.95 ± 0.07, 1-sample Wilcoxon signed-rank test against 1, *W*_71_ = -638, *P* = 0.073, *n* = 72 neurons). (e-f) Scatterplots of the A vs. B lap correlation against the performance from all sessions and animals during learning (e) and recall (f). A linear regression fit revealed a strong inverse relationship between the A-B correlation maps and performance for the learning cohort, but not for the recall cohort (**Learning**: linear fit, *R* = -0.69, \*\*\**P* = 2.17e-07, n = 44 sessions from 6 mice; **Recall**: linear fit, *R* = -0.32, *P* = 0.058, n = 35 sessions from 5 mice). Error bars indicate **95% ±C**.**I. around median** for normalized correlations scores and mean ± s.e.m. for performance fraction.

To further link the differential activity of CA1 neurons to task performance, we used population vector decoding (see Methods) to read out both the position and the context (A or B trial) of the mice. For each training session, we used the first half of the session to train a linear decoder to predict absolute position as well as trial context (A vs B trials) in the second half (Fig. 6a-d). The decoder’s performance was lower in early training sessions, when behavioral performance was low, compared with later sessions, when both behavioral performance and decoding accuracy increased (Fig. 6e-f). Notably, decoder accuracy in distinguishing context A from B was significantly correlated (Fig. 6g) with behavioral performance across all mice, while absolute error in decoding position was not (Fig. 6h). Thus, we observe that improvement in performance during training is closely tracked by the ability of the population of neurons to discriminate between the two contexts. Closer inspection revealed that the decoder improved its ability to distinguish trial context the most around position 110 cm in context A, immediately before the reward zone for that context (Fig. 6i). Context accuracy was also lower near the end of trials, in both contexts, but this did not improve with experience.

**Fig. 6:**
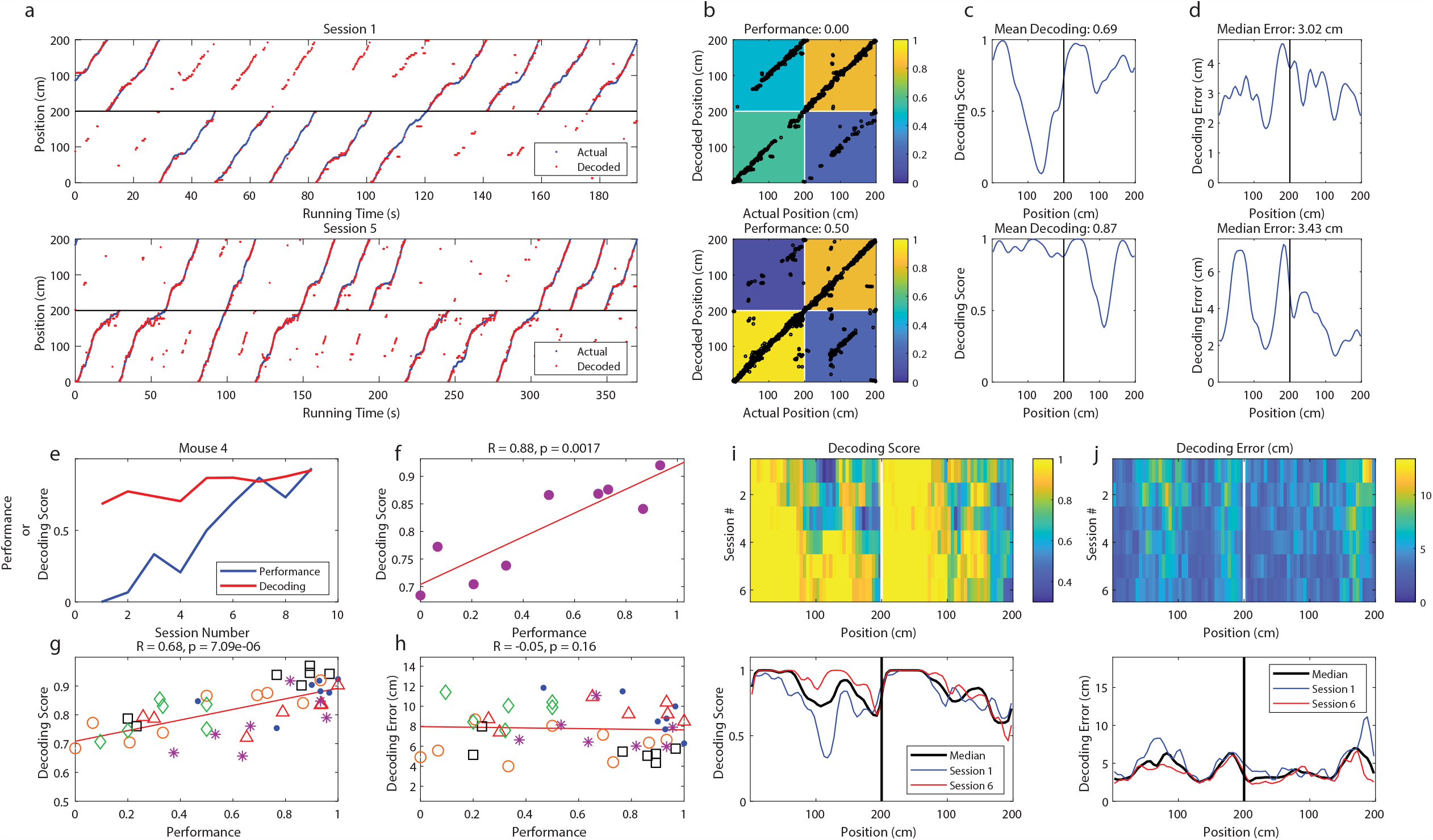
Place maps incorporate information about trial type during task learning and accurately predict both location and trial during recall. (a) A population vector decoder accurately predicts the position of an animal on the track during both the initial training session (top) and late training session (bottom). Plotted are the decoded positions (red) against the actual track position (blue) of the mouse as a function of time. On the late training session, the decoder for this animal additionally predicts the trial the animal is on with high accuracy. Throughout the figure, positions 0-200 are represented twice, once each for A and B trials. Data from this mouse only are used in panels b-d. (b) Confusion matrices quantifying decoding accuracy demonstrate improved accuracy in identifying trial type associated with improved performance. Each black point represents the actual position and trial of the animal plotted against the decoder’s prediction. Each matrix cell represents the number of decoded points falling into each quadrant divided by the total data points in each trial type. (c) Average trial decoding score (proportion of data points correctly classified) as a function of position in early sessions (top) and late sessions (bottom). The majority of misclassifications occurred early in training in the middle and late stages of the track on A trials. (d) The median position decoding error across the track on both trials did not substantially change across learning. (e) Example plot showing that as the performance of a single mouse (ID: 4) increases with subsequent training sessions, the accuracy of the population decoder also increases. (f) Same data as in (e) with performance plotted against decoding score, revealing a strong positive relationship between task performance and decoder score (**Correlation score significance**: two-sided one-sample t-test, *t*_7_ = 4.948 \*\**P* = 0.002, n = 9 sessions from mouse 4). (g) Cumulative analysis across all training sessions revealed a strong positive relationship between performance and decoding accuracy (**Correlation between performance and decoding score in learning cohort**: two-way ANOVA, *R* = 0.68, effect of mouse *F*_5,31_ = 0.338, *P* = 0.338, effect of performance, *F*_1,31_ = 29.014, \*\*\**P* < 0.001, effect of interaction, *F*_5,31_ = 0.806, *P* = 0.554, n = 43 session from 6 mice). Each point denotes a single training session and each type of mark a different animal. (**h**) In contrast, no relationship was observed between position decoding error and task performance (**Correlation between performance and decoding score in recall cohort:** two-way ANOVA, *R* = -0.05, effect of mouse, *F*_5,31_ = 3.057, \**P* = 0.023, effect of performance *F*_1,31_ = 2.026, *P* = 0.164, effect of interaction, *F*_5,31_ = 1.569, *P* = 0.197, n = 43 session from 5 mice). (i) Colormap showing the improvement in the position-trial decoding accuracy as function of training session. Each row denotes the median decoding score across all 6 animals. The decoding score increases progressively forward along the track during each training session, prominently observed on A trials. Bottom graph plots decoding score for sessions 1 and 6, as well as the median across all sessions. (j) An equivalent plot of the decoding position error averaged across all animals does not reveal such change during training.

Errors in absolute position also tended to be higher near the end of trials in both contexts, as well as immediately after reward zone B experienced in context A (Fig. 6j). These regions of decreased decoder accuracy correspond to parts of trials with the most uncertainty in the neural population, which may ultimately contribute to behavioral errors. The performance of the decoder thus suggests that spatial maps in CA1 do not require *de novo* mapping of the spatial environment to integrate trial-specific information, but rather that contextual information becomes integrated with pre-existing maps during learning.

## Discussion

In this study, we developed a one-dimensional, head-fixed, odor-cued navigation behavior to examine hippocampal spatial map dynamics during learning and recall, similar to freely moving behavioral tasks such as the continuous alternation T-maze^17^. Our task provides two distinct advantages over previous freely moving studies. First, we trained animals to learn the task contingency on the exact same belt. This way, we can ensure hippocampal representation of the task is due to operant learning of a behavior with defined episodic, spatial and contextual components, rather than the learning of other strategies such as dead reckoning. Second, we capitalized on ultra-stable two-photon calcium imaging of the same population of transgenically expressed GCaMP+ neurons to longitudinally track the emergence and remote retrieval of place maps under this behavioral paradigm. Our results fuel evidence for two long-postulated features of hippocampal network dynamics: that existing place maps can reconfigure in response to an associational learning rule despite a constant physical environment, and that such task selective place maps persist long after learning is achieved.

Our data, where we observed emergence of cells with place fields exclusively in behavioral context A, or B, shows that the hippocampus generated task-selective representations of space during learning that rapidly remapped as a function of odor-cued behavioral contingency. Imaging of spatial activity during our task in well-trained mice revealed a complex set of coding mechanisms for conjunctive representation of both location and behavioral context. In cells with fields in both behavioral contexts, A and B, we for the first time observed the calcium analog of two well-described remapping properties of place cells recorded in freely-moving behavior – rate and global remapping – whose activity is attributed to changes in context^13, 33^ and physical environment^13, 21^, respectively. Given the structure of our task, where the physical environment remained constant while the odor-cued behavioral context changed, we were surprised to see a relatively smaller fraction of activity remapping place cells (analogous to rate remapping) as compared to global and more complex remapping cells. There are two possible explanations for this discrepancy. First, despite enhanced sensitivity of the GCaMP6f Ca^2+^indicator^27^ used in our imaging, calcium as a proxy for neuronal activity may not resolve smaller changes in activity rates (or spike modulation) resulting in an underestimation of rate remapping place cells. We attempt to address this by using duration and AUC of calcium events as remapping metrics rather than amplitude, as they better reflect changes in the bursting activity of neurons. Alternatively, the high proportion of global remapping cells is observed because animals use different trial specific spatial reference frames to navigate toward reward zones.

Our study, along with others, shows that hippocampal “place cells” can modulate their activity rate or switch their place field tuning in different environments, and even within the same environment given changes in task demands or goal locations^13, 17^-^19, 22, 23, 34^. At the ensemble level, the tuning and density of trial selective place cells are structured according to the episodic and spatial salience associated with the trial context. Over-representation of these place fields not only occurs selectively around the trial-respective reward locations, but in an episodically relevant goal directed manner. For example, in A-trials, animals must traverse and deliberately withhold licking at the B-reward zone before reaching the targeted A-reward zone. Following this, place cell density is high in the cue-sampling, B- *and* A-reward zones. However, field density for B-selective place cells drop soon after crossing reward goal location B (Zone I), signifying the spatio-contextual irrelevance of the rest of the belt for the given trial.

In spite of the prominence of place cells (and other feature-selective neurons^15-17, 35-37^) in the hippocampus, their relationship to learning and execution of learnt behaviors remain controversial. To directly quantify the behavioral correlates for emergence of spatial selectivity to temporally structured rule-based learning we used neural decoder. This allowed us to go beyond the standard prediction of position and context (trial type) information during learnt behavior^7, 38, 39^. Here, we observed a very strong correlation between task performance and the accuracy with which the hippocampal map could predict the location and trial type the animal was in. Very early in learning, absolute location on the track could be decoded. Only later in learning could behavioral context be decoded, mirroring rate of learning. Otherwise stated, the transformation of spatial maps was not associated with a significant loss of spatial information during learning, but rather with the accuracy with which the network could predict the current trial type at any given location. Although we hypothesized that the substrate of this learning would be an increase in the proportion of cells uniquely representing A or B behavioral contexts, we instead found that the proportion of cells tuned to each behavioral context did not change significantly. Rather, learning appeared to be driven by increased distance between place fields of cells jointly representing context A and B.

Lastly, in contrast to previous studies that show a high degree of CA1 place cell instability across time, representational drift was greatly reduced across remote retrieval sessions, likely because animals were task-engaged rather than randomly foraging.

This stabilization of place maps following learning resolves a debate in the field: how can the day-to-day instability of place cells reported in recent studies be reconciled with the hypothesis that they serve as the substrate for stable long-term episodic memories. Our findings expand on previous *in vivo* electrophysiological recordings in CA1, which first reported the increased stability of place fields during memory-guided, attention-dependent behavior over a 6 hour interval^5^. In contrast, rapid turnover of spatial map activity in CA1 was observed in imaging experiments during less structured behaviors such as random foraging^7, 23^, goal-oriented learning^23^, and non-operantly rewarded spatial context switching^8, 23^. Our results show such instability of place map activity can be significantly reduced when the map is embedded within an operant rule-based learning regimented by contextual, episodic and spatial feature selection. Our result bolsters the importance of behavioral state on the stability of hippocampal representations, alongside a growing body of work binding spatial and non-spatial coding in the hippocampus with learning and attention. For example, learning of a fear association with a particular environment induces remapping and stabilization of place cells in the long-term^40^. Olfactory and visuo-spatial representations show enhanced stability and fidelity for recall with attentional demands^41^. Further, the same ‘odor cells’ in dorsal CA1 are reactivated across days following learning of an olfactory delayed working-memory task^42^. Beyond the hippocampus, emergence of stable and sparse representations with learning was observed in the motor cortex^43^.

What are the cellular and circuit mechanisms driving the task-specific place cell dynamics we observe? The olfactory cue context and navigational demands of our task likely relies on recruiting interactions with lateral entorhinal cortex (LEC)^44, 45^ and medial entorhinal cortex (MEC)^46-55^, but perhaps during distinct task phases. LEC lesions impairs rate remapping in CA3 place^33^, and may be involved in driving context dependent remapping^52, 56^ during the learning phases of our task. On the other hand, MEC lesions or input manipulations disrupt place cell precision and stability^57, 58^ as well as place memory although only partially^59^, implicating a role for MEC in the stabilization of spatial activity following learning. Coordinated activity and integration of entorhinal cortex and CA3 inputs upon CA1 pyramidal neurons can result in dendritic spikes^60, 61^. These dendritic spikes have been implicated in context discrimination behavior^61^ and context-dependent place cell formation and remapping^62^, potentially through recruitment of non-Hebbian plasticity mechanisms like input timing dependent plasticity (ITDP) and behavioral time-scale dependent plasticity (BTSP), during the learning phase of the task. Whereas, potential mechanisms driving stabilized ensemble coding in the long term following learning include Hebbian plasticity rules that involve theta modulated post synaptic burst firing^63-65^, and experience dependent strengthening of coincident spatially tuned synaptic inputs^66^.

Consistent with previous findings^66, 67^, our trial-by-trial remapping occurs on very fast timescales, well below the temporal regime of typical plasticity mechanisms. Such a fast context dependent switch could be supported by specific input gating or gain control through modulation of inhibitory ^67-69^ and disinhibitory circuits^61^, and are worth exploring in the context of our observed behaviorally modulated activity. In terms of stabilization of subsets of place maps during the remote recall phases of the task, higher order prefrontal cortical (PFC)^70-72^, and subcortical neuromodulatory inputs^5, 73, 74^ may be at play. Reactivation and stabilization^75^ of place cell sequences has also been attributed to highly synchronous sharp-wave ripple (SPW-Rs) activity in CA1. Such, SWRs erupt during immobility or slow wave sleep in a strongly correlated but time compressed fashion to prior task performance^22, 76^, future navigation decisions^36, 77^, and perhaps are at play during the memory consolidation phase of our task.

In conclusion, our data shows that the hippocampus rapidly generated task-selective representations of space during learning. Moreover, the emergent ensembles used both simple and complex remapping of their activity for alternating between spatial representations on different trials. Interestingly, while behaviorally-influenced representations of space emerged early on, these maps continued to evolve towards progressively more dissimilar cross-trial representations. These were inversely correlated to increasing animal performance. While much work remains to uncover the possible cellular and circuit mechanisms driving the experience dependent emergence and stabilization of place cell ensembles, this novel behavioral paradigm provides a rich substrate to study flexibility and stability of place maps in episodic and context-dependent manners.

## Author contributions

RZ and JB conceived the project, designed the experiments and wrote the paper, RZ performed the experiments and data analysis, JJM built the neural decoder and performed analysis for figure 5 and 6.

## Acknowledgments

This work was supported by an NIH BRAIN INITIATIVE 1R01NS109994, an NIH 1R01NS109362-01, McKnight Scholar Award in Neuroscience, Klingenstein-Simons Fellowship Award in Neuroscience, Alfred P.Sloan Research Fellowship, Whitehall Research Grant, American Epilepsy Society Junior Investigator Award, Blas Frangione Young Investigator Research Grant, New York University Whitehead Fellowship for Junior Faculty in Biomedical and Biological Sciences and the Leon Levy Foundation Award to JB. RZ was supported by an NYU Grossman School of Medicine MSTP NIH 5T32 GM007308 grant; JJM was supported by an HHMI extension grant to Dmitri Chkolvskii. We are indebted to György Buszáki, Dmitry Rinberg, Vincent Robert, Rachel Swanson, and Olesia Bilash for helpful discussions on the study and comments on previous versions of the manuscript and Cara Johnson for proofreading the document.

## Methods

### Mice

Experiments were performed with 4-12 month old adult male mice on a C57BL/6J background transgenically expressing GCaMP6f from the *Thy1* locus (GP5.5 JAX strain #024276)^78^. All experiments were approved by the Institutional Animal Care and Use Committee at New York University Medical Center.

### Hippocampal Window and Headpost Implantation

Mice were implanted with a circular imaging window (3.0 mm x 1.5 mm [diameter x height]) centered at 2.3 AP and 1.5 ML over the left dorsal-intermediate hippocampus surrounded by a modified 3-D printed headpost^79^ for head fixation. Imaging cannulas were made by attaching a 3-mm diameter coverslip (64-0720 Warner) to a stainless steel cylindrical cannula using optical UV curing optical adhesive (NOA-61, Norland products)^23^.

### *In Vivo* Two-Photon Imaging

Imaging was performed using a two-photon 8-kHz resonant scanner (Ultima, Bruker) with a 16x, 0.8 NA water-immersion objective (Nikon). Excitation was performed at 920 nm with an 80 MHz pulsed laser (Mai Tai DeepSee, Spectra Physics). GCaMP6f emission fluorescence was collected with a GaAsP photomultiplier tube (7422P-40, Hamamatsu) following red and green channel separation with a filter cube consisting of a dichroic mirror (T565lpxr, Chroma Technology) and filters (green, ET510/80m-2p; red, ET605/70m-2p, Chroma Technology). Images were acquired at a 30 Hz frame rate, 512×512 pixel resolution, and 1.5x digital zoom corresponding to a field size of 555 µm x 555 µm.

### Behavior

*Behavioral apparatus*. Mice ran on a custom-built treadmill track where the belt consisted of 3 ∼65 cm long distinct fabrics (macro-textures) enriched with 4 micro-textures (5 cm regions consisting of 5 ‘dice’ arranged flattened aluminum foil spheres, 4 crossed hook-and-loop strips, zig-zag glue pattern, and strip of woven material). The position of the mouse was measured using an optical rotary encoder (S5-720, US Digital). Lap onset and micro-texture crossings were detected by reading associated RFID tags with an RFID reader mounted below the animal (ID-20LA, SparkFun Electronics). Behavior tones of 4 kHz,10 kHz, and white noise were pre-recorded and played using an mp3 player (MP3 Player Shield, DEV-12660, SparkFun Electronics). The audio signal from each channel was then amplified (PAM8302, Adafruit) and played though a pair of speakers (25-1719S, Tang Band) located on the sides of the fixation platform. Licking of the animal was registered via a blunt-tipped steel canula coupled to a capacitive touch sensor (SEN-12041, SparkFun Electronics). Behavioral programs were controlled with an Arduino Mega 2560 microcontroller. All behavioral data was acquired at a sampling rate of 10 kHz with a data acquisition board (PCI-6052E, National Instruments) synchronized to the time of frame acquisition.

#### Olfactometer

A custom-built olfactometer was used for delivering fixed-concentration, spatially restricted odors to the mouse (Supplemental Fig.1). Briefly, during non-odor zone navigation, animals were exposed to a constant background flow of air mixed with pure mineral oil (BP26291, Fisher Scientific) at a flow rate of 1 L/min. In the immediate 50 cm prior to odor zone entry, an odor charge program was initiated by closing the normally open (NO) inlet/outlet valves (225T021, NResearch Inc.) along the flow path through the mineral oil vial (M), while opening the flow path through either odor A or B vials by activating the respective normally closed (NC) inlet/outlet valves (225T011, NResearch Inc.). The odor-charged air was routed to an exhaust port during this period through a 3-way final valve (SH360T041, NResearch Inc.), while 1 L/min of air continued to be delivered. Charging was performed in order to ensure that a consistent steady-state concentration of odor was reached along the pre-delivery flow path (before the final valve) and to minimize latency of odor delivery to the animal. Upon entering the odor zone, the final valve was triggered via an RFID tag to switch routing of the odor-charged air from exhaust to the animal’s snout. Upon reaching the end of the odor zone, an RFID tag triggered the closure of either A or B path valves and opening of the background mineral oil air path. A constant vacuum of 1L/min above the odor delivery port ensured scavenging of residual odors. 10% dilutions of pentyl acetate (Sigma-Aldrich, 109584) and (+)-α-Pinene (Sigma-Aldrich, P45680) in mineral oil were used as odor A and B, respectively. A photoionization detector (200B: miniPID Dispersion Sensor, Aurora Scientific) was used to verify steady-state odor concentration delivery prior to each imaging session. A steady-state odor onset latency (baseline to steady-state) of ∼125 ms and off latency of ∼75 ms (steady-state to baseline) was measured. Fresh dilutions of odors were prepared daily.

#### Random foraging

Following recovery from surgery (3-5 days), mice were water deprived and habituated to handling and head-fixation to behavioral apparatus. Water-deprived mice were then trained to operantly lick and receive 5% sucrose water rewards in regularly spaced reward zones along a ∼196 cm linear track consisting of fabrics and textures described above. Training began with 20 regularly distributed reward zones followed by a program of progressively fewer and more randomly distributed reward zones over 2 weeks. Access to ∼1 μL sucrose droplets began immediately after entry into a reward zone and terminated following either an exit from the reward zone (20 cm initial length), a time-out period (7 s initial duration) that had elapsed since entry, or once a maximum number of collected rewards had been reached (10 reward initial limit). Sucrose rewards were delivered on alternate licks. Training was considered complete once mice ran at a rate of ∼1 lap/min in search of 3 random reward zones per lap, each defined by a 10 cm length, 3 s time-out period, and 5 droplet collection limit. Animals were given a total of 1 mL of water daily.

#### Odor-cued spatial navigation task

Following successful training on random reward foraging, animals were introduced onto a structured training regimen that consisted of alternating blocks of A and B trial laps. On the first day of training, animals were placed on an alternating sequence of A and B laps for 10 trials to familiarize the animal with the two types of trials. Thereafter, a regimen of 5 sequential A and 5 sequential B laps (5A5B) was presented in an alternating block pattern which progressed to an alternating block of 3 A and 3 B laps (3A3B) and finally to randomized lap (random) presentation.

Each rewarded lap was signaled by a 0.5 s 4 kHz tone immediately prior to lap start. If the animal reached the start of the lap prior to the 0.5 s elapsing, the tone would stop playing. The odor was delivered across the initial 20 cm segment of the lap. Delivery of the trial-associated reward was restricted to the 10 cm reward zone, a 3 s collection time, and a maximum of 10 rewards. Following >∼80% task performance, incorrect behavior was punished by time-out laps. When the animal licked in either the anticipatory (10 cm prior to reward zone) or reward zone not associated with the current trial, a 0.5 s 10 kHz tone was played, signaling to the animal an incorrect choice. On the following time-out lap, a 0.5 s white noise was played prior to lap entry, neither odor A nor B was delivered, and no reward was available.

### Image processing and signal extraction

#### Motion correction and ROI segmentation

Imaging time-series data was corrected for motion artifacts by using the NoRMCorre non-rigid motion correction algorithm implemented in MATLAB^80^. The first imaging session in each longitudinal imaging series was used as the template against which all future sessions were motion corrected.

Segmentation of somatic regions of interest (ROIs) was performed using a constrained non-negative matrix factorization (CNMF) approach implemented in MATLAB as part of the CaImAn software package^30, 81^. Non-somatic and low-quality components were manually discarded using a custom graphical interface.

#### Matching components across sessions

Individually identified somatic ROIs in each session were matched across sessions by using the *register_multisession*.*py* function as part of the CaImAn Python package. Matched components across all sessions were subsequently visualized and poorly matching or mismatching components were discarded. Discarding of component matches was blind to the calcium signal associated with a component on any given session.

#### Relative fluorescence change

(ΔF/F) The signal baseline (*F*_0 *baseline*_) was calculated for each ROI by taking the fluorescence signal and calculating its 50^th^ percentile (median) value at each timepoint within a sliding 15s time window using the *prctfilt*.*m* function as part of CaImAn. The same procedure was used to extract the background signal from the background component (*F*_0 *background*_). The ΔF/F was then calculated as:

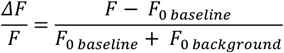

The resultant ΔF/F signal was subsequently smoothed using an exponential filter with *τ* = 0.2*s* to reduce photon shot noise from signal acquisition^82^.

#### Calcium event detection

Significant calcium events were identified using an algorithm previously used in the analysis of two-photon, CA1 hippocampal imaging data^23, 83, 84^. Briefly, for any given ΔF/F calcium trace, deflections from the baseline value due to acquisition noise and/or motion along the dorsoventral (z) axis should occur with equal frequency in both the positive and negative directions. Based on this assumption, the false-positive rate can be calculated for each putative event and an amplitude and duration threshold can be defined such that an event’s false-positive event rate does not exceed 5% (rate at which positive events occur with at least 20-fold higher probability than negative events). Using this approach, we identified initial putative events by detecting consecutive imaging frames whose onset occurred at 2 s.d. above the mean and whose offset occurred at 0.5 s.d. below the mean. All events within a session were classified according to their amplitude (in 0.5 sigma bins) and duration (in 250 ms bins). We calculated the false-positive rate for each amplitude-duration bin as the ratio of negative to positive events in that bin. Only positive events from bins with a false-positive ratio of less than 5% were included in the analysis.

To further improve the sensitivity of event detection, initially detected events were masked on the original fluorescence signal, the *F*_0 *baseline*_ was recalculated, and events were redetected on the updated ΔF/F signal. Two iterations of event-masked baseline recalculation were performed. Events that lasted less than 1 s were excluded from subsequent analysis.

### Data analysis

#### Definition of run epochs

As described previously^23, 83^, we defined running epochs as consecutive frames during which the mouse was moving forward with a minimum peak speed of 5 cm/s for at least 1 s in duration. Neighboring run epochs separated by less than 0.5 s were merged. All other epochs were defined as no run.

#### Selection of place cells

*Spatial information* We identified spatially tuned cells (place cells) by computing their spatial information content relative to an empirically generated shuffle distribution as described previously^23, 83^. The spatial information content was defined as^85^:

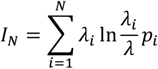

Where *λ*_*i*_ is the transient rate and *p*_*i*_ is fraction of running time spent in the *i*th spatial bin, *λ* is the overall transient rate, and *N* is the number of bins. The transient rate was defined as the ratio of the bin count of running-related transient onsets smoothed with a Gaussian kernel (σ = 3 bins) to the spatial bin occupancy time. We computed *I*_*N*_ for *N* = 2,4,5,10,20,25,50,100 bins. To create shuffle distributions for each of the *N* spatial bins, we randomly reassigned the transient onset times within the running-related epochs 1,000 times and recomputed the spatial information content for each reassignment 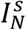, where *s* is the index of the shuffle. To approximately account for the bias associated with spatial binning in the calculation of the spatial information content, we subtracted the mean of the shuffled null distribution from each *N*-binned estimate to obtain the adjusted *I*_*N*_ values:

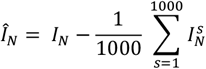

We then obtained a single estimate of the spatial information content for each neuron by taking the maximum of the adjusted information values 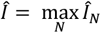 for the true transient onset times and the shuffled onset times 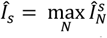. The spatial tuning p-value was defined as the fraction of shuffle values s for which Î exceeded Î s. Neurons with a spatial tuning p-value < 0.05 were defined as place cells. *Tuning specificity* We calculated the spatial tuning vector for each cell as described previously^23^ using the formula 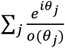, where is the binned position of the mouse (*N* = 200 bins, 1 bin ∼ 1 cm) at the onset time of the *j*^th^ run-epoch transient, and *o*_*j*_ is the occupancy of the animal at position *θ*_*j*_, i.e. the fraction of running frames that the animal spent at position *θ*_*j*_. Calculation of the spatial tuning vector was restricted to only run epochs as defined above. The tuning specificity was defined as the magnitude of the spatial tuning vector. Statistical significance of the tuning specificity for each cell was determined by first generating a null tuning distribution by shuffling the transient onset times within the run-epoch frames and then computing the tuning specificity from each shuffle. The shuffle was performed 1,000 times for each cell and the p-value was defined as the fraction of the null distribution that exceeded the cell’s actual tuning specificity.

#### Activity rate

We calculated the activity rate as the cumulative area under the ΔF/F traces (AUC), from event onset to offset, of all significant calcium events in either run or no-run epochs and divided this sum by the amount of time the animal spent in respective epochs.

#### Place fields

To define the width of place fields, we first calculated the rate map for each neuron by dividing the run-epoch event count in each spatial bin by the bin occupancy for *N* = 100 bins and then smoothed using a Gaussian kernel (σ = 3). To define spatially significant fields, we then fit each local maximum in the rate map with a Gaussian and defined the width as the distance between the locations where each fitted curve was at 20% of its peak value. Putative overlapping fields were merged into single fields. Only fields with a minimum of 5 significant events on distinct laps were included in the analysis.

#### Spatial tuning curves (STC)

The tuning curves were defined as the ratio of a Gaussian-smoothed (σ = 3) count of significant run-epoch calcium events in each bin (*N* = 100) to the run-epoch occupancy. Each neuron’s tuning curve was normalized to its maximum activity across both trial types. For visualization purposes, tuning curves were smoothed again with a Gaussian kernel (σ = 3).

### Task-selective and remapping place cell selection criteria

#### Task-selective neurons

Selective neurons were initially chosen as those which were spatially tuned by either the spatial information or tuning specificity criterion in one set of trials and by neither in the other set of trials. Only neurons that had at least 5 in-field, run-epoch calcium events on distinct laps and those in which the animal was in a run epoch 80% of the time of the equivalent spatial bin range of these calcium events on the other trial laps were included in analysis.

#### Common and global remapping neurons

To determine which neurons globally remapped, we performed a Pearson correlation of their rate maps (*N* = 100 bins) between correct A and B laps. Correlation was performed only between spatial bins with non-zero values in either trial. Neurons that had a positive, statistically significant (p-value < 0.05) correlation score were classified as common neurons (their spatial maps were similar), while neurons with non-significant scores (p >= 0.05) or significant negative scores (maps which are either dissimilar or anti-correlated) were classified as globally remapping. We verified that the distributions of correlation scores against their p-values for all common and global neurons separated into two distinct classes (Supplementary Fig. 6). Only neurons that were tuned according to tuning specificity were used in the analysis. All cells were required to have a single place field on each set of trials and at least 5 significant calcium events on distinct laps in their place fields. Additionally, for globally remapping neurons, animals must have been in a run epoch at least 80% of the time within the equivalent range of calcium onset bins of the other trials on at least 6 laps (to ensure that the animal was in a run epoch on both lap types).

#### Activity remapping neurons

Among the neurons that were selected as common, we examined the area under the curve (AUC) of in-field calcium events to determine whether there was a significant variation in activity associated with trial type. Given that animal speed contributes to CA1 place cell firing activity^86-88^, we performed a 2-way ANOVA test to determine the effect of task trial type and speed on the AUC of calcium events. Neurons that had a trial type effect p-value < 0.05 were classified as activity remapping. We further confirmed that this category was distinct from the common population by calculating the difference over sum ratio of the peak of the average of calcium transients between the in-field events of correct A trials and correct B trials (Supplementary Fig. 8).

#### Partial remapping neurons

Neurons with partially remapping fields were selected as those which met either the spatial information or tuning specificity criterion in both trial types and had 2 place fields in one type of trials, whereas only 1 place field in the other. For a place field to be considered common across trial types, the distance between the centroid of the place fields between trial types must have been less than the 95^th^ percentile value of the distribution of the place field centroids for the common neurons. No threshold was set on the distance of the remapping field centroid from the common field. As with global remapping neurons, all place fields were required to have at least 5 significant in-field calcium events on distinct laps. For the partial field, the animal must have been in a run epoch at least 80% of the time within the equivalent range of calcium onset bins of the other trial type on at least 6 laps.

#### Population vector correlation

The normalized spatial tuning curves across 100 spatial bins were assembled for all neurons into a 2D matrix where the rows represented neuron indexes and columns the activity of all the neurons in each spatial bin. Thus, each column represented the population activity of all neurons at a particular bin. For similarity analysis, the population vector in each column was Pearson correlated against a different trial set or imaging session. The mean of the correlation scores from all bins was the population vector correlation score.

#### Tuning vector correlation

Between spatially tuned neurons on any two sessions, the spatial tuning vectors across 100 bins were Pearson correlated for each neuron and the mean of all correlated neurons was the tuning vector correlation score.

### Population Vector Decoding

To demonstrate the relationship between behavioral performance and tuning fidelity of our recorded neurons, we performed population vector decoding (Fig. 6). A separate decoder was constructed for each session for each mouse. For a given session, template tuning curves for each cell were constructed in a similar manner as described above, only using data when mice were running. Briefly, we divided the 200 cm track into 40 bins each for A and B trials (80 bins total), counted the number of calcium events in each bin, and smoothed with a Gaussian smoothing kernel with *σ* = 5 cm, then divided by the total time spent in each bin. Data from the first half of the session was used to define the template. Time-varying rate vectors for each cell were constructed using data from the second half of the session using 250 ms bins, smoothed with a Gaussian smoothing kernel with sigma *σ* = 250 ms. For each time point in the second half of the session, the decoded position was the position corresponding to the highest correlation with the template matrix.

The performance of the decoder was quantified using two measures: “Decoding Score” and “Decoding Error.” Decoding score (Fig. 6c, e, f, g, i) was defined as the proportion of data points that were correctly classified as belonging to A or B trials. Decoding Error (Fig. 6d, h, j) was defined as the mean absolute distance between the decoded and actual position when ignoring trial type. Distance was defined in a circular manner such that positions 0 and 200 were at the same point.

#### Spatial raster plots

Lap-by-lap raster plots were made by taking the mean run-epoch ΔF/F value from 100 spatial bins for each place cell.

#### Statistics

Statistical analysis of calcium data was done using Matlab R2020a (Mathworks).

#### Software

All analysis was done using custom-written scripts in MATLAB R2020a (Mathworks). Scripts are available on request from R.Z. and J.B.

## References

1. Tulving, E. Elements of Episodic Memory (Oxford University Press, 1985).

2. Tulving, E. Episodic Memory: From Mind to Brain. Annual Review of Psychology 53, 1–25 (2002).

3. O’Keefe, J. The hippocampus as a cognitive map / John O’Keefe and Lynn Nadel (Clarendon Press ; Oxford University Press, Oxford : New York, 1978).

4. O’Keefe, J. & Dostrovsky, J. The hippocampus as a spatial map. Preliminary evidence from unit activity in the freely-moving rat. Brain Research 34, 171–175 (1971).

5. Kentros, C.G., et al. Increased Attention to Spatial Context Increases Both Place Field Stability and Spatial Memory. Neuron 42, 283–295 (2004).

6. Kentros, C. Hippocampal place cells: The “where” of episodic memory? Hippocampus 16, 743–754 (2006).

7. Ziv, Y., et al. Long-term dynamics of CA1 hippocampal place codes. Nature Neuroscience 16, 264 (2013).

8. Hainmueller, T. & Bartos, M. Parallel emergence of stable and dynamic memory engrams in the hippocampus. Nature 558, 292–296 (2018).

9. Scoville, W.B. & Milner, B. Loss of recent memory after bilateral hippocampal lesions. Journal of Neurology, Neurosurgery & Psychiatry 20, 11–21 (1957).

10. Tulving, E. & Markowitsch, H.J. Episodic and declarative memory: Role of the hippocampus. Hippocampus 8, 198–204 (1998).

11. Rolls, E.T. A computational theory of episodic memory formation in the hippocampus. Behavioural Brain Research 215, 180–196 (2010).

12. Markus, E.J., et al. Interactions between location and task affect the spatial and directional firing of hippocampal neurons. The Journal of Neuroscience 15, 7079 (1995).

13. Leutgeb, S., et al. Independent Codes for Spatial and Episodic Memory in Hippocampal Neuronal Ensembles. Science 309, 619 (2005).

14. Muller, R.U. & Kubie, J.L. The effects of changes in the environment on the spatial firing of hippocampal complex-spike cells. The Journal of Neuroscience 7, 1951 (1987).

15. Komorowski, R.W., Manns, J.R. & Eichenbaum, H. Robust Conjunctive Item– Place Coding by Hippocampal Neurons Parallels Learning What Happens Where. The Journal of Neuroscience 29, 9918 (2009).

16. Wood, E.R., Dudchenko, P.A. & Eichenbaum, H. The global record of memory in hippocampal neuronal activity. Nature 397, 613–616 (1999).

17. Wood, E.R., Dudchenko, P.A., Robitsek, R.J. & Eichenbaum, H. Hippocampal Neurons Encode Information about Different Types of Memory Episodes Occurring in the Same Location. Neuron 27, 623–633 (2000).

18. Ferbinteanu, J. & Shapiro, M.L. Prospective and Retrospective Memory Coding in the Hippocampus. Neuron 40, 1227–1239 (2003).

19. Frank, L.M., Brown, E.N. & Wilson, M. Trajectory Encoding in the Hippocampus and Entorhinal Cortex. Neuron 27, 169–178 (2000).

20. Hayashi, Y. NMDA Receptor-Dependent Dynamics of Hippocampal Place Cell Ensembles. The Journal of Neuroscience 39, 5173 (2019).

21. Leutgeb, S., et al. Distinct ensemble codes in hippocampal areas CA3 and CA1. Science 305, 1295–1298 (2004).

22. Dupret, D., O’Neill, J., Pleydell-Bouverie, B. & Csicsvari, J. The reorganization and reactivation of hippocampal maps predict spatial memory performance. Nature Neuroscience 13, 995–1002 (2010).

23. Danielson, N.B., et al. Sublayer-Specific Coding Dynamics during Spatial Navigation and Learning in Hippocampal Area CA1. Neuron 91, 652–665 (2016).

24. Whitlock, J.R., Heynen, A.J., Shuler, M.G. & Bear, M.F. Learning induces long-term potentiation in the hippocampus. science 313, 1093–1097 (2006).

25. Pastalkova, E., et al. Storage of spatial information by the maintenance mechanism of LTP. science 313, 1141–1144 (2006).

26. Skaggs, W.E., McNaughton, B.L., Wilson, M.A. & Barnes, C.A. Theta phase precession in hippocampal neuronal populations and the compression of temporal sequences. Hippocampus 6, 149–172 (1996).

27. Chen, T.-W., et al. Ultrasensitive fluorescent proteins for imaging neuronal activity. Nature 499, 295–300 (2013).

28. Muller, R. A Quarter of a Century of Place Cells. Neuron 17, 813–822 (1996).

29. Muller, R.U., et al. Spatial firing correlates of neurons in the hippocampal formation of freely moving rats. in Brain and space. 296–333 (Oxford University Press, New York, NY, US, 1991).

30. Giovannucci, A., et al. CaImAn an open source tool for scalable calcium imaging data analysis. eLife 8, e38173 (2019).

31. Ainge, J.A., van der Meer, M.A.A., Langston, R.F. & Wood, E.R. Exploring the role of context-dependent hippocampal activity in spatial alternation behavior. Hippocampus 17, 988–1002 (2007).

32. Goshen, I., et al. Dynamics of Retrieval Strategies for Remote Memories. Cell 147, 678–689 (2011).

33. Lu, L., et al. Impaired hippocampal rate coding after lesions of the lateral entorhinal cortex. Nature neuroscience 16, 1085 (2013).

34. Leutgeb, J.K., et al. Progressive Transformation of Hippocampal Neuronal Representations in “Morphed” Environments. Neuron 48, 345–358 (2005).

35. Aronov, D., Nevers, R. & Tank, D.W. Mapping of a non-spatial dimension by the hippocampal–entorhinal circuit. Nature 543, 719–722 (2017).

36. Pastalkova, E., Itskov, V., Amarasingham, A. & Buzsáki, G. Internally Generated Cell Assembly Sequences in the Rat Hippocampus. 321, 1322–1327 (2008).

37. MacDonald, Christopher J., Lepage, Kyle Q., Eden, Uri T. & Eichenbaum, H. Hippocampal “Time Cells” Bridge the Gap in Memory for Discontiguous Events. Neuron 71, 737–749 (2011).

38. Rubin, A., Geva, N., Sheintuch, L. & Ziv, Y. Hippocampal ensemble dynamics timestamp events in long-term memory. eLife 4, e12247 (2015).

39. Brown, E.N., et al. A Statistical Paradigm for Neural Spike Train Decoding Applied to Position Prediction from Ensemble Firing Patterns of Rat Hippocampal Place Cells. 18, 7411–7425 (1998).

40. Wang, M.E., et al. Long-Term Stabilization of Place Cell Remapping Produced by a Fearful Experience. The Journal of Neuroscience 32, 15802 (2012).

41. Muzzio, I.A., et al. Attention Enhances the Retrieval and Stability of Visuospatial and Olfactory Representations in the Dorsal Hippocampus. PLOS Biology 7, e1000140 (2009).

42. Taxidis, J., et al. Emergence of stable sensory and dynamic temporal representations in the hippocampus during working memory. 474510 (2018).

43. Peters, A.J., Chen, S.X. & Komiyama, T. Emergence of reproducible spatiotemporal activity during motor learning. Nature 510, 263–267 (2014).

44. Xu, W. & Wilson, D.A. Odor-evoked activity in the mouse lateral entorhinal cortex. Neuroscience 223, 12–20 (2012).

45. Leitner, F.C., et al. Spatially segregated feedforward and feedback neurons support differential odor processing in the lateral entorhinal cortex. Nature Neuroscience 19, 935–944 (2016).

46. Miao, C., et al. Hippocampal Remapping after Partial Inactivation of the Medial Entorhinal Cortex. Neuron 88, 590–603 (2015).

47. Sargolini, F., et al. Conjunctive Representation of Position, Direction, and Velocity in Entorhinal Cortex. Science 312, 758–762 (2006).

48. Fyhn, M., et al. Spatial Representation in the Entorhinal Cortex. Science 305, 1258–1264 (2004).

49. Hafting, T., et al. Microstructure of a spatial map in the entorhinal cortex. Nature 436, 801–806 (2005).

50. Stensola, H., et al. The entorhinal grid map is discretized. Nature 492, 72–78 (2012).

51. Deshmukh, S. & Knierim, J. Representation of Non-Spatial and Spatial Information in the Lateral Entorhinal Cortex. 5 (2011).

52. Neunuebel, J.P., Yoganarasimha, D., Rao, G. & Knierim, J.J. Conflicts between Local and Global Spatial Frameworks Dissociate Neural Representations of the Lateral and Medial Entorhinal Cortex. 33, 9246–9258 (2013).

53. Knierim, J.J., Neunuebel, J.P. & Deshmukh, S.S. Functional correlates of the lateral and medial entorhinal cortex: objects, path integration and local-global reference frames. Philos Trans R Soc Lond B Biol Sci 369, 20130369–20130369 (2013).

54. Deshmukh, S.S., Yoganarasimha, D., Voicu, H. & Knierim, J.J. Theta Modulation in the Medial and the Lateral Entorhinal Cortices. Journal of Neurophysiology 104, 994–1006 (2010).

55. Kropff, E., Carmichael, J.E., Moser, M.-B. & Moser, E.I. Speed cells in the medial entorhinal cortex. Nature 523, 419–424 (2015).

56. Wang, C., et al. Egocentric coding of external items in the lateral entorhinal cortex. Science 362, 945–949 (2018).

57. Jacob, P.-Y., et al. Medial entorhinal cortex lesions induce degradation of CA1 place cell firing stability when self-motion information is used. Brain and Neuroscience Advances 4, 2398212820953004 (2020).

58. Bittner, K.C., et al. Conjunctive input processing drives feature selectivity in hippocampal CA1 neurons. Nature Neuroscience 18, 1133–1142 (2015).

59. Hales, Jena B., et al. Medial Entorhinal Cortex Lesions Only Partially Disrupt Hippocampal Place Cells and Hippocampus-Dependent Place Memory. Cell Reports 9, 893–901 (2014).

60. Kamondi, A., Acsády, L. & Buzsáki, G. Dendritic Spikes Are Enhanced by Cooperative Network Activity in the Intact Hippocampus. 18, 3919–3928 (1998).

61. Basu, J., et al. Gating of hippocampal activity, plasticity, and memory by entorhinal cortex long-range inhibition. Science 351, aaa5694 (2016).

62. Bittner, K.C., et al. Behavioral time scale synaptic plasticity underlies CA1 place fields. 357, 1033–1036 (2017).

63. Epsztein, J., Lee, A.K., Chorev, E. & Brecht, M. Impact of Spikelets on Hippocampal CA1 Pyramidal Cell Activity During Spatial Exploration. Science 327, 474–477 (2010).

64. Epsztein, J., Brecht, M. & Lee, Albert K. Intracellular Determinants of Hippocampal CA1 Place and Silent Cell Activity in a Novel Environment. Neuron 70, 109–120 (2011).

65. Lee, D., Lin, B.-J. & Lee, A.K. Hippocampal Place Fields Emerge upon Single-Cell Manipulation of Excitability During Behavior. Science 337, 849–853 (2012).

66. Cohen, J.D., Bolstad, M. & Lee, A.K. Experience-dependent shaping of hippocampal CA1 intracellular activity in novel and familiar environments. eLife 6, e23040 (2017).

67. Sheffield, M.E.J., Adoff, M.D. & Dombeck, D.A. Increased Prevalence of Calcium Transients across the Dendritic Arbor during Place Field Formation. Neuron 96, 490-504.e495 (2017).

68. Lovett-Barron, M., et al. Dendritic Inhibition in the Hippocampus Supports Fear Learning. Science 343, 857–863 (2014).

69. Turi, G.F., et al. Vasoactive Intestinal Polypeptide-Expressing Interneurons in the Hippocampus Support Goal-Oriented Spatial Learning. Neuron 101, 1150-1165.e1158 (2019).

70. Preston, Alison R. & Eichenbaum, H. Interplay of Hippocampus and Prefrontal Cortex in Memory. Current Biology 23, R764–R773 (2013).

71. Spellman, T., et al. Hippocampal–prefrontal input supports spatial encoding in working memory. Nature 522, 309 (2015).

72. Rajasethupathy, P., et al. Projections from neocortex mediate top-down control of memory retrieval. Nature 526, 653–659 (2015).

73. Solari, N. & Hangya, B. Cholinergic modulation of spatial learning, memory and navigation. Eur J Neurosci 48, 2199–2230 (2018).

74. Kaufman, A.M., Geiller, T. & Losonczy, A. A Role for the Locus Coeruleus in Hippocampal CA1 Place Cell Reorganization during Spatial Reward Learning. Neuron 105, 1018-1026.e1014 (2020).

75. Roux, L., et al. Sharp wave ripples during learning stabilize the hippocampal spatial map. Nature Neuroscience 20, 845 (2017).

76. Jadhav, S.P., Kemere, C., German, P.W. & Frank, L.M. Awake Hippocampal Sharp-Wave Ripples Support Spatial Memory. 336, 1454–1458 (2012).

77. Ito, H.T., et al. A prefrontal–thalamo–hippocampal circuit for goal-directed spatial navigation. Nature 522, 50 (2015).

78. Dana, H., et al. Thy1-GCaMP6 Transgenic Mice for Neuronal Population Imaging In Vivo. PLOS ONE 9, e108697 (2014).

79. Osborne, J.E. & Dudman, J.T. RIVETS: A Mechanical System for In Vivo and In Vitro Electrophysiology and Imaging. PLOS ONE 9, e89007 (2014).

80. Pnevmatikakis, E.A. & Giovannucci, A. NoRMCorre: An online algorithm for piecewise rigid motion correction of calcium imaging data. Journal of Neuroscience Methods 291, 83–94 (2017).

81. Pnevmatikakis, Eftychios A., et al. Simultaneous Denoising, Deconvolution, and Demixing of Calcium Imaging Data. Neuron 89, 285–299 (2016).

82. Jia, H., Rochefort, N.L., Chen, X. & Konnerth, A. In vivo two-photon imaging of sensory-evoked dendritic calcium signals in cortical neurons. Nature Protocols 6, 28–35 (2011).

83. Zaremba, J.D., et al. Impaired hippocampal place cell dynamics in a mouse model of the 22q11.2 deletion. Nature Neuroscience 20, 1612–1623 (2017).

84. Dombeck, D.A., et al. Imaging Large-Scale Neural Activity with Cellular Resolution in Awake, Mobile Mice. Neuron 56, 43–57 (2007).

85. Skaggs, W.E., McNaughton, B.L. & Gothard, K.M. An Information-Theoretic Approach to Deciphering the Hippocampal Code. in Advances in Neural Information Processing Systems 5, [NIPS Conference] 1030–1037 (Morgan Kaufmann Publishers Inc., 1993).

86. McNaughton, B., Barnes, C.A. & O’Keefe, J. The contributions of position, direction, and velocity to single unit activity in the hippocampus of freely-moving rats. Experimental brain research 52, 41–49 (1983).

87. Ekstrom, A., Meltzer, J., McNaughton, B. & Barnes, C.A. NMDA receptor antagonism blocks experience-dependent expansion of hippocampal “place fields”. Neuron 31, 631–638 (2001).

88. Czurkó, A., Hirase, H., Csicsvari, J. & Buzsáki, G. Sustained activation of hippocampal pyramidal cells by ‘space clamping’in a running wheel. European Journal of Neuroscience 11, 344–352 (1999).

