## Supplemental Figures for "Behaviorally emergent hippocampal place maps remain stable during memory recall"

**Extended Data Fig. 1: Olfactometer design and operation.**

Schematic illustrating the position- and trial-controlled delivery of odors to the mouse. Clean, in-house air is supplied to a pressure regulator which reduces incoming pressure to the olfactometer to 10 psi. During continuous delivery of air containing mineral-oil (background air), incoming air is routed through three flowmeters. Air through the first flowmeter is routed through a vial containing mineral oil (M path) at a rate of 0.1 L/min through normally open valves and is diluted by merging with air coming through the second flowmeter at a rate of 0.9 L/min at the manifold. Air from the manifold is sent through the final valve after which it arrives at the mouse at a rate of 1 L/min. The remainder of the incoming air is routed through the third flowmeter at a rate of 1 L/min through the final valve and into an exhaust outlet. Prior to odor delivery to the mouse, an odor charge cycle is initiated to achieve steady-state flow through the final valve. Upon charge cycle initiation, the input and output valves through the mineral oil vial are closed, while either odor A or B input and output valves are opened and mix with the clean air at the manifold. Simultaneously, the final valve is switched and routes the odor charged air to the exhaust, while delivering non-odorized air the mouse. When the mouse enters the odor zone, the final valve is switched again, and the charged odor air passing though the valve is delivered to the mouse at low latency. At the end of the odor zone, the odor is discharged from the delivery line by opening the mineral oil valves (M path) and closing either the A or B (A/B path) input/output valves. A constant vacuum is applied near the nose cone where the odor (or background air) is delivered at a negative flow rate of 1 L/min, equal to the final odor flow at the mouse's snout, to clear residual odors. NO – normally-open valve, NC – normally-closed valve.

**Extended Data Fig. 2: Peri-reward zone speed across training stages.**

Analysis of speed in the area immediately prior to reward zone entry and within the zone across progressive training stages revealed a consistent decline in mean speed within trial-appropriate reward zones. (a) Animal speed in the pre- and post- A reward zone area across training stages on A laps. A statistically significant drop in speed was observed on all training stages with no difference in speed observed during random foraging (**Pre- vs. post- A reward zone speed on A laps - RF**: two-tailed paired *t*-test, $t_3 = 0.557$, $P = 0.617$, n = 4 mice; **5A5B**: two-tailed paired *t*-test, $t_3 = 6.748$, $^{**}P = 0.007$, n = 4 mice; **3A3B**: two-tailed paired *t*-test, $t_3 = 8.716$, $^{**}P = 0.003$, n = 4 mice; **Random**: two-tailed paired *t*-test, $t_3 = 13.699$, $^{***}P < 0.001$, n = 4 mice). (b) In contrast, no significant trend was observed in the A reward zone on B trial laps with no significant difference on the last training stage (**Pre- vs. post- A reward zone speed on B laps – 5A5B**: two-tailed paired *t*-test, $t_3 = 3.60$, $^{*}P = 0.037$, n = 4 mice; **3A3B**: two-tailed paired *t*-test, $t_3 = 1.163$, $P = 0.329$, n = 4 mice; **Random**: two-tailed paired *t*-test, $t_3 = 2.701$, $P = 0.074$, n = 4 mice). Similarly, a significant speed decrease in reward zone B was observed on B trials during the training stages (c) with no such trend observed on A trial laps (d; **Pre- vs. post- B reward zone speed on B laps - RF**: two-tailed paired *t*-test, $t_3 = 1.037$, $P = 0.376$, n = 4 mice; **5A5B**: two-tailed paired *t*-test, $t_3 = 4.967$, $^{*}P = 0.016$, n

= 4 mice; **3A3B**: two-tailed paired *t*-test, $t_3$ = 5.337, *$P$ = 0.013, n = 4 mice; **Random**: two-tailed paired *t*-test, $t_3$ = 9.377, **$P$ = 0.003, n = 4 mice; **Pre- vs. post- B reward zone speed on A laps – 5A5B**: two-tailed paired *t*-test, $t_3$ = 1.632, $P$ = 0.201, n = 4 mice; **3A3B**: two-tailed paired *t*-test, $t_3$ = 3.453, *$P$ = 0.041, n = 4 mice; **Random**: two-tailed paired *t*-test, $t_3$ = 0.140, $P$ = 0.898, n = 4 mice). Speed was calculated over a 2 second interval along the length of track immediately prior to zone entry and 2 second interval following zone entry across all laps. Error bars represent mean ± s.e.m. Paired t-tests were used to calculated significance.

**Extended Data Fig. 3: Lap speed of each animal along track during learned behavior.**

(a) For animals used to analyze the trial-selective and remapping properties of place cells in Figs. 2 and 3, all n = 10 mice (11 FOVs) showed similar spatial bin speeds across the track with a characteristic decline in speed in the trial-appropriate peri-reward zones. The mean animal speed was ~15-20 cm/s. Dotted lines represent the spatial bin where the A (blue) or B (red) trial reward zone begins. Solid lines represent mean spatial bin speed across the respective trials laps and shaded area is the s.e.m. in that spatial bin. The track was subdivided into 100 spatial bins for analysis. (b) There was no substantial difference between animal speed between A and B trials (less than 5 cm/s difference) except near the trial-specific reward zones where the animal was expected to slow down. Data shown as mean ± s.e.m.

**Extended Data Fig. 4: Place field properties and in-field speed of task-selective and non-selective place cells.**

(a) The mean speed of mice during crossing of task-selective place fields remained similar regardless of which trial lap the animal was in. Each point represents a task-selective neuron with its mean in-field speed (cm/s) in A vs. B laps for A-selective (left) neurons and B-selective neurons (right). (b) A- and B- selective place cells had a significant but small mean speed difference between trial types (**A-B place field speed difference in A-selective neurons**: two-tailed 1-sample Wilcoxon signed-rank test against 0, $W_{586}$ = -65174, ***$P$ < 0.001, $n$ = 587 neurons; **A-B place field speed difference in B-selective neurons**: two-tailed 1-sample Wilcoxon signed-rank test against 0, $W_{465}$ = 44165, ***$P$ < 0.001, $n$ = 466 neurons). Place field properties of task-selective place cells did not significantly differ compared to task nonselective place cells. (c) Most task-selective place cells had single place fields on both A and B trials with a higher fraction of B-selective place cells having single place fields compared to place fields on B trials for task non-selective place cells. A-selective place cells were less likely to have two place fields relative to task nonselective place cells on A trials (**A- vs. B-selective single place field count**: two-tailed paired Wilcoxon signed-rank test, $W_{10}$ = -14, $P$ = 0.577, $n$ = 11 FOV from 10 mice, Holm-Sidak correction; **A-selective vs. A&B-A single place field count**: two-tailed paired Wilcoxon signed-rank test, $W_{10}$ = 66, **$P$ = 0.003, $n$ = 11 FOV from 10 mice, Holm-Sidak correction; **A-selective vs. A&B-B single place field count**: two-tailed paired Wilcoxon signed-rank test, $W_{10}$ = 66, **$P$ =

0.003, $n$ = 11 FOV from 10 mice, Holm-Sidak correction; **A- vs. B-selective double place field count**: two-tailed paired Wilcoxon signed-rank test, $W_{10}$ = -10, $P$ = 0.7, $n$ = 11 FOV from 10 mice, Holm-Sidak correction; **A-selective vs. A&B-A double place field count**: two-tailed paired Wilcoxon signed-rank test, $W_{10}$ = -66, **$P$ = 0.003, $n$ = 11 FOV from 10 mice, Holm-Sidak correction; **A-selective vs. A&B-B double place field count**: two-tailed paired Wilcoxon signed-rank test, $W_{10}$ = -66, **$P$ = 0.003, $n$ = 11 FOV from 10 mice, Holm-Sidak correction; **A- vs. B-selective triple place field count**: two-tailed paired Wilcoxon signed-rank test, $W_{10}$ = 28, *$P$ = 0.031, $n$ = 11 FOV from 10 mice, Holm-Sidak correction; **A-selective vs. A&B-A triple place field count**: two-tailed paired Wilcoxon signed-rank test, $W_{10}$ = -37, $P$ = 0.064, $n$ = 11 FOV from 10 mice, Holm-Sidak correction; **A-selective vs. A&B-B triple place field count**: two-tailed paired Wilcoxon signed-rank test, $W_{10}$ = -64, **$P$ = 0.006, $n$ = 11 FOV from 10 mice, Holm-Sidak correction). (d) The distribution of place field widths did not differ between selective and non-selective place cells (**A- vs. B- selective place field width**: 2-sample Kolmogorov-Smirnov test, $D_{695, 542}$ = 0.09, $P$ = 0.069, $n$ = 695 vs. 542 neurons, 11 FOV from 10 mice, Holm-Sidak correction; **A- vs. A&B-A- selective place field width**: 2-sample Kolmogorov-Smirnov test, $D_{695, 3655}$ = 0.03, $P$ = 0.765, $n$ = 695 vs. 3655 neurons, 11 FOV from 10 mice, Holm-Sidak correction; **A- vs. A&B-B selective place field width**: 2-sample Kolmogorov-Smirnov test, $D_{695, 3533}$ = 0.05, $P$ = 0.286, $n$ = 695 vs. 3533 neurons, 11 FOV from 10 mice, Holm-Sidak correction; **B- vs. A&B-A selective place field width**: 2-sample Kolmogorov-Smirnov test, $D_{542, 3655}$ = 0.1, **$P$ = 0.002, $n$ = 542 vs. 3655 neurons, 11 FOV from 10 mice, Holm-Sidak correction; **B- vs. A&B-B selective place field width**: 2-sample Kolmogorov-Smirnov test, $D_{542, 3533}$ = 0.04, $P$ = 0.765, $n$ = 542 vs. 3533 neurons, 11 FOV from 10 mice, Holm-Sidak correction; **A&B-A vs. A&B-B selective place field width**: 2-sample Kolmogorov-Smirnov test, $D_{3655, 3533}$ = 0.06, ***$P$ < 0.001, $n$ = 3655 vs. 3533 neurons, 11 FOV from 10 mice, Holm-Sidak correction). All data shown as mean ± s.e.m.

**Extended Data Fig. 5: Common and globally remapping place cells are classified by the correlation between their spatial tuning curves.**

Globally remapping place cells display significant shifts in their place field tuning curves between trials. (a) A non-remapping place cell with a common place field that does not shift between lap trials and demonstrates a positive spatial tuning curve correlation value. (b) A remapping place cell with a shifting place field between trials shows a negative spatial tuning curve correlation indicating remapping (c) Scatterplot of the Pearson's correlation value (r) of spatial tuning curves against the negative logarithm of the associated p values for each place cell having a significant place field in both A and B trials. Place cells that had either negative and/or a correlation p-value < 0.05 were considered global remapping place cells (magenta) while the remaining cells were classified as common (purple). Cross mark on the right side of the scatter plot indicates common place cell shown in (a), while cross mark on the left side shows global remapping place cell in (b). Dashed horizonal line is the negative logarithm of cutoff p-value = 0.05 and vertical dashed line represents a correlation score of 0. The list of all correlation scores and corresponding p-values can be found in Table 1.

**Extended Data Fig. 6: Distribution of distances between trial-specific place fields of globally remapping place cells.**

Global remapping place cells had place fields that remapped across the entire length of the track with the majority of place fields showing mean centroid shifts of less than 30 cm. (a) Imaging fields of view with three examples of globally remapping place cells categorized according to the distance between their place field centroids on A vs. B trials. (b) Scatterplot of the place field centroids for all globally remapping neurons on A vs. B trials. Place cells with place fields on A laps tend to shift to earlier locations on the lap on B trials, particularly in Zone II of the track (Fig. 3f). Dashed red line indicates location of the start of reward zone B, while dashed blue line indicates the start of reward zone B. (c) Distribution of place field centroid differences between A and B laps. The centroid difference is skewed toward shorter distances with a median of ~31 cm. (d) Examples of three globally remapping neurons with increasing place field centroid differences. Magnified FOVs correspond to the neurons labeled in (a) with calcium traces, polar event plots, and spatial tuning curves, respectively, shown on the right.

**Extended Data Fig. 7: Activity discrimination index validates sub-classification of activity remapping neurons from common neuron pool.**

The activity discrimination index validates the presence of a distinct, activity-remapping subpopulation of common place cells that share the same place field on A and B trials, but differ by their trial-specific in-field place activity. (a) A cumulative density plot showing a significant difference in the distribution of activity discrimination scores between in-field place field activity on A vs. B trials calculated according to the formula shown below (**Common vs global remapping activity index score**: 2-sample Kolmogorov-Smirnov test, $D_{700, 64} = 0.24$, $^{**}P = 0.001$, $n = 700$ vs. 64 neurons, 11 FOV from 10 mice). (b) Mean in-field calcium transients aligned to the onset time of the transients for activity remapping place cells which show higher mean in-field activity on A trials (top) and those which higher in-field mean activity on B trials (bottom). Neurons are shown according to increase activity index score from left to right. (c) The overall activity rate during run epochs was not different for activity remapping neurons between A and B trials suggesting that the activity difference is specific to the place field (**Activity rate in A vs. B laps during RUN for all activity remapping neurons**: two-tailed paired Wilcoxon signed-rank test, $W_{63} = 470$, $P = 0.116$, $n = 64$ neurons).

**Extended Data Fig. 8: Example of manual parsing of automatically matched pyramidal neuron soma ROIs across sessions**

Example FOVs from an animal displayed in Fig. 4 showing spatial component matching of soma across non-consecutive imaging sessions. Both upper and lower panels show spatial components of neurons that were matched using an automatic script as part of the CaImAn calcium imaging analysis software. The top panel shows neurons which were manual selected from automatically matched components for analysis on session 1 (cyan) and session 3 (yellow) with their merge show in magenta. Notice the high degree of component overlap for the selected components between session 1 and 3. Bottom panel shows the matched components that were excluded from analysis due to partial or incomplete spatial overlap or mismatch.

**Extended Data Fig. 9: Equivalent characterization of spatial stability and learning-dependent spatial tuning decorrelation using spatial information criterion**

Using the spatial information (S.I.) tuning selection criterion, the results of learning-dependent decorrelation and stability of recall maps were similar to those we observed when using the tuning specificity (T.S.) spatial tuning criterion. (a) The fraction of place cells tuned in each trial did not show any significant trend during learning (left) or recall (right; **Fraction of A-trial tuned place cells during learning - S.I.**: one-way RM mixed effects analysis, effect of training day, $F_{6, 21.93} = 2.34$, $P = 0.067$, $n = 6$ mice; **Day 1 vs. Day 6 A trial learning**: two-tailed paired *t*-test, $t_4 = -0.94$, $P = 0.475$, $n = 5$ mice, Holm-Sidak correction; **Day 1 vs. Day 7 A trial learning**: two-tailed paired *t*-test, $t_3 = -1.33$, $P = 0.475$, $n = 4$ mice, Holm-Sidak correction; **Fraction of B-trial tuned place cells during learning - S.I.**: one-way RM mixed effects analysis, effect of training day, $F_{6, 27} = 1.55$, $P = 0.201$, $n = 6$ mice; **Day 1 vs. Day 6 B trial learning**: two-tailed paired *t*-test, $t_4 = -0.21$, $P = 0.847$, $n = 5$ mice, Holm-Sidak correction; **Day 1 vs. Day 7 B trial learning**: two-tailed paired *t*-test, $t_3 = -0.93$, $P = 0.664$, $n = 4$ mice, Holm-Sidak correction; **Fraction of A-trial tuned place cells during recall - S.I.**: one-way RM mixed effects analysis, effect of training day, $F_{6, 24} = 0.39$, $P = 0.88$, $n = 5$ mice; **Day 1 vs. Day 6 A trial recall**: two-tailed paired *t*-test, $t_4 = -0.63$, $P = 0.674$, $n = 5$ mice, Holm-Sidak correction; **Day 1 vs. Day 7 A trial recall**: two-tailed paired *t*-test, $t_4 = -0.88$, $P = 0.674$, $n = 5$ mice, Holm-Sidak correction; **Fraction of B-trial tuned place cells during recall - S.I.**: one-way RM mixed effects analysis, effect of training day, $F_{6, 24} = 0.72$, $P = 0.637$, $n = 5$ mice; **Day 1 vs. Day 6 B trial recall**: two-tailed paired *t*-test, $t_4 = -2.05$, $P = 0.207$, $n = 5$ mice, Holm-Sidak correction; **Day 1 vs. Day 7 B trial recall**: two-tailed paired *t*-test, $t_4 = 0.06$, $P = 0.955$, $n = 5$ mice, Holm-Sidak correction). (b) The correlation of spatial tuning curves for S.I.-tuned neurons relative to the first day of imaging similarly showed a significantly steeper decline during learning compared to recall on A and B trials (**TC correlation on A trials**: two-way RM mixed effects analysis, effect of time, $F_{3, 22.23} = 57.28$, $***P < 0.001$, effect of behavior, $F_{1, 8.93} = 8.08$, $*P = 0.019$, interaction between time and behavior, $F_{3, 22.23} = 3.46$, $*P = 0.034$, $n = 6$ learn cohort, 5 recall cohort mice; **Day 2 vs. Day 6 A trial learning**: two-tailed paired *t*-test, $t_3 = 18.46$, $***P < 0.001$, $n = 4$ mice, Holm-Sidak correction; **Day 2 vs. Day 7 A trial learning**: two-tailed paired *t*-test, $t_2 = 8.1$, $*P = 0.015$, $n = 3$ mice, Holm-Sidak correction; **TC correlation on B trials**: two-way RM mixed effects analysis, effect of time, $F_{3, 22.64} = 57.54$, $***P < 0.001$, effect of behavior, $F_{1, 9.14} = 17.28$, $**P = 0.002$, interaction between time and behavior, $F_{3, 22.64} = 2.02$, $P = 0.14$, $n = 6$ learn cohort, 5 recall cohort mice; **Day 2 vs. Day 6 B trial learning**: two-tailed paired *t*-test, $t_3 = 3.47$, $*P = 0.04$, $n = 4$ mice, Holm-Sidak correction; **Day 2 vs. Day 7 B trial learning**: two-tailed paired *t*-test, $t_2 = 8.55$, $*P$

= 0.027, $n$ = 3 mice, Holm-Sidak correction; **Day 2 vs. Day 6 A trial recall**: two-tailed paired $t$-test, $t_4 = 4.19$, $^*P = 0.014$, $n$ = 5 mice, Holm-Sidak correction; **Day 2 vs. Day 7 A trial recall**: two-tailed paired $t$-test, $t_4 = 5.89$, $^{**}P = 0.008$, $n$ = 5 mice, Holm-Sidak correction; **Day 2 vs. Day 6 B trial recall**: two-tailed paired $t$-test, $t_4 = 19.07$, $^{***}P <$ 0.001, $n$ = 5 mice, Holm-Sidak correction; **Day 2 vs. Day 7 B trial recall**: two-tailed paired $t$-test, $t_4 = 11.71$, $^{***}P < 0.001$, $n$ = 5 mice, Holm-Sidak correction; **Day 6 A trials learning vs. recall**: two-tailed unpaired $t$-test, $t_8 = -4.27$, $^{**}P = 0.005$, $n$ = 5 learn cohort, 5 recall cohort mice, Holm-Sidak correction; **Day 7 A trials learning vs. recall**: two-tailed unpaired $t$-test, $t_7 = -3.6$, $^{**}P = 0.009$, $n$ = 4 learn cohort, 5 recall cohort mice, Holm-Sidak correction; **Day 6 B trials learning vs. recall**: two-tailed unpaired $t$-test, $t_8 = -2.8$, $^*P = 0.023$, $n$ = 5 learn cohort, 5 recall cohort mice, Holm-Sidak correction; **Day 7 B trials learning vs. recall**: two-tailed unpaired $t$-test, $t_7 = -5.21$, $^{**}P = 0.002$, $n$ = 4 learn cohort, 5 recall cohort mice, Holm-Sidak correction). (c) The same pattern of stability emerged when S.I. tuned place cells were used for the neighboring day correlation analysis during learning and recall. Learning and recall plots split for ease of visualization (**Neighboring session TC correlation on A trials**: two-way RM mixed effects analysis, effect of time, $F_{2,\,14.33} = 2.93$, $P = 0.086$, effect of behavior, $F_{1,\,9.2} = 3.7$, $P = 0.086$, interaction between time and behavior, $F_{2,\,14.33} = 5.47$, $^*P = 0.017$, $n$ = 6 learn cohort, 5 recall cohort mice; **Neighboring session TC correlation on B trials**: two-way RM mixed effects analysis, effect of time, $F_{2,\,13.92} = 0.81$, $P = 0.466$, effect of behavior, $F_{1,\,8.61} = 4.55$, $P = 0.063$, interaction between time and behavior, $F_{2,\,13.92} = 6.72$, $^{**}P =$ 0.009, $n$ = 6 learn cohort, 5 recall cohort mice; **Days 1 vs. 2 Vs. Day 6 vs. 7 A trials learn**: two-tailed paired $t$-test, $t_2 = -15.6$, $^{**}P = 0.004$, $n$ = 3 mice; **Days 1 vs. 2 Vs. Day 6 vs. 7 B trials learn**: two-tailed paired $t$-test, $t_2 = -6.36$, $^*P = 0.024$, $n$ = 3 mice; **Days 1 vs. 2 Vs. Day 6 vs. 7 A trials recall**: two-tailed paired $t$-test, $t_4 = 0.32$, $P = 0.764$, $n$ = 5 mice; **Days 1 vs. 2 Vs. Day 6 vs. 7 B trials recall**: two-tailed paired $t$-test, $t_4 = 1.4$, $P =$ 0.234, $n$ = 5 mice). (d) The mean centroid difference between place fields relative to day 1 of imaging increased more during learning compared to recall on A and B laps (**Mean centroid difference relative to Day 1 A trials**: two-way RM mixed effects analysis, effect of time, $F_{3,\,22.44} = 28.61$, $^{***}P < 0.001$, effect of behavior, $F_{1,\,8.9} = 2.6$, $P = 0.141$, interaction between time and behavior, $F_{3,\,22.44} = 2.67$, $P = 0.072$, $n$ = 6 learn cohort, 5 recall cohort mice; **Mean centroid difference relative to Day 1 B trials**: two-way RM mixed effects analysis, effect of time, $F_{3,\,24.47} = 17.03$, $^{***}P < 0.001$, effect of behavior, $F_{1,\,9.49} = 11.83$, $^{**}P = 0.007$, interaction between time and behavior, $F_{3,\,24.47} = 0.06$, $P =$ 0.981, $n$ = 6 learn cohort, 5 recall cohort mice; **Day 5 A trials learning vs. recall**: two-tailed unpaired $t$-test, $t_8 = 2.36$, $P = 0.09$, $n$ = 5 learn cohort, 5 recall cohort mice, Holm-Sidak correction; **Day 5 B trials learning vs. recall**: two-tailed unpaired $t$-test, $t_8 = 2.17$, $P = 0.119$, $n$ = 5 learn cohort, 5 recall cohort mice, Holm-Sidak correction). (e) The decorrelation between A&B tuned place cells was also inversely correlated to the animals' performance during training (left) in contrast to recall session (right; **Day one normalized A vs. B lap correlation scores for matching neurons during learning**: Kruskal-Wallis test, $H_5 = 105.83$, $^{***}P < 0.001$, $n$ = 1967 neurons from 6 mice; **Day 2 learn**: two-tailed 1-sample Wilcoxon signed-rank test against 1, $W_{316} = -10531$, $^{**}P =$

0.001, $n$ = 317 neurons; **Day 7 learn**: two-tailed 1-sample Wilcoxon signed-rank test against 1, $W_{236}$ = -16113, ***$P$ < 0.001, $n$ = 237 neurons; **Day one normalized A vs. B lap correlation scores for matching neurons during recall**: Kruskal-Wallis test, $H_5$ = 4.34, $P$ = 0.502, $n$ = 1577 neurons from 5 mice; **Day 2 recall**: two-tailed 1-sample Wilcoxon signed-rank test against 1, $W_{338}$ = 228, $P$ = 0.95, $n$ = 339 neurons; **Day 7 recall**: two-tailed 1-sample Wilcoxon signed-rank test against 1, $W_{228}$ = -2699, $P$ = 0.179, $n$ = 229 neurons). All data represented as mean ± s.e.m.

**Extended Data Fig. 10: Long-term stability of place maps during recall.**

Place maps remained stable beyond the one-week imaging interval with a slow decay in map correlation relative to first imaging day on subsequent weeks. (a) The relative fraction of place cells tuned to either trial type remained stable across weeks with no trend in relative distribution using either spatial information (S.I.; left) and tuning specificity (T.S.; right) to select spatial tuned place cells (**Fraction of A-trial tuned place cells during long-term recall - S.I.**: one-way RM mixed effects analysis, effect of training day, $F_{5, 7.07}$ = 2.01, $P$ = 0.193, $n$ = 3 mice; **Day 1 vs. Day 25 A trial recall**: two-tailed paired $t$-test, $t_2$ = -0.64, $P$ = 0.588, $n$ = 3 mice, Holm-Sidak correction; **Day 1 vs. Day 30 A trial recall**: two-tailed paired $t$-test, $t_2$ = -2.8, $P$ = 0.203, $n$ = 3 mice, Holm-Sidak correction; **Fraction of B-trial tuned place cells during long-term recall - S.I.**: one-way RM mixed effects analysis, effect of training day, $F_{5, 7.05}$ = 0.47, $P$ = 0.789, $n$ = 3 mice; **Day 1 vs. Day 25 B trial recall**: two-tailed paired $t$-test, $t_2$ = -0.1, $P$ = 0.93, $n$ = 3 mice, Holm-Sidak correction; **Day 1 vs. Day 30 B trial recall**: two-tailed paired $t$-test, $t_2$ = -0.69, $P$ = 0.808, $n$ = 3 mice, Holm-Sidak correction; **Fraction of A-trial tuned place cells during long-term recall - T.S.**: one-way RM mixed effects analysis, effect of training day, $F_{5, 7.13}$ = 1.83, $P$ = 0.224, $n$ = 3 mice; **Day 1 vs. Day 25 A trial recall**: two-tailed paired $t$-test, $t_2$ = -0.52, $P$ = 0.693, $n$ = 3 mice, Holm-Sidak correction; **Day 1 vs. Day 30 A trial recall**: two-tailed paired $t$-test, $t_2$ = -0.94, $P$ = 0.693, $n$ = 3 mice, Holm-Sidak correction; **Fraction of B-trial tuned place cells during long-term recall - T.S.**: one-way RM mixed effects analysis, effect of training day, $F_{5, 7.54}$ = 0.34, $P$ = 0.877, $n$ = 3 mice; **Day 1 vs. Day 25 B trial recall**: two-tailed paired $t$-test, $t_2$ = 0.09, $P$ = 0.939, $n$ = 3 mice, Holm-Sidak correction; **Day 1 vs. Day 30 B trial recall**: two-tailed paired $t$-test, $t_2$ = -3.32, $P$ = 0.154, $n$ = 3 mice, Holm-Sidak correction). (b) The performance of all imaged animals (n=3) remained stable at or near 100% on all long-term recall sessions. (c) The stability of spatial maps dropped significantly on sixth day imaging session, but remained at stable correlation score ~0.4-0.5 on sessions thereafter for all neurons (left) and those significantly tuned to space on using S.I. or T.S criteria (right; **PV correlation during long-term recall relative to day 1 on A trials**: one-way RM mixed effects analysis, effect of training day, $F_{4, 5.08}$ = 11.39, **$P$ = 0.01, $n$ = 3 mice; **Day 1 vs. Day 25 A trial recall**: two-tailed paired $t$-test, $t_1$ = 3.82, $P$ = 0.3, $n$ = 2 mice, Holm-Sidak correction; **Day 1 vs. Day 30 A trial recall**: two-tailed paired $t$-test, $t_1$ = 3.36, $P$ = 0.3, $n$ = 2 mice, Holm-Sidak correction; **PV correlation during long-term recall relative to day 1 on B trials**: one-way RM mixed effects analysis, effect of training day, $F_{4, 5.01}$ = 78.75, ***$P$ < 0.001, $n$ = 3 mice; **Day 1 vs. Day 25 B trial recall**: two-tailed paired $t$-test,

$t_1$ = 11.33, *P* = 0.061, *n* = 2 mice, Holm-Sidak correction; **Day 1 vs. Day 30 B trial recall**: two-tailed paired *t*-test, $t_1$ = 20.54, *P* = 0.061, *n* = 2 mice, Holm-Sidak correction; **TC correlation (S.I.) during long-term recall relative to day 1 on A trials**: one-way RM mixed effects analysis, effect of training day, $F_{4, 5.09}$ = 20.19, **P = 0.003, *n* = 3 mice; **Day 1 vs. Day 25 A trial recall**: two-tailed paired *t*-test, $t_1$ = 5.5, *P* = 0.157, *n* = 2 mice, Holm-Sidak correction; **Day 1 vs. Day 30 A trial recall**: two-tailed paired *t*-test, $t_1$ = 7.74, *P* = 0.157, *n* = 2 mice, Holm-Sidak correction; **TC (S.I.) correlation during long-term recall relative to day 1 on B trials**: one-way RM mixed effects analysis, effect of training day, $F_{4, 5.04}$ = 43.78, ***P < 0.001, *n* = 3 mice; **Day 1 vs. Day 25 B trial recall**: two-tailed paired *t*-test, $t_1$ = 8.95, *P* = 0.088, *n* = 2 mice, Holm-Sidak correction; **Day 1 vs. Day 30 B trial recall**: two-tailed paired *t*-test, $t_1$ = 14.17, *P* = 0.088, *n* = 2 mice, Holm-Sidak correction; **TC correlation (T.S.) during long-term recall relative to day 1 on A trials**: one-way RM mixed effects analysis, effect of training day, $F_{4, 5.21}$ = 15.82, **P = 0.004, *n* = 3 mice; **Day 1 vs. Day 25 A trial recall**: two-tailed paired *t*-test, $t_1$ = 3.32, *P* = 0.186, *n* = 2 mice, Holm-Sidak correction; **Day 1 vs. Day 30 A trial recall**: two-tailed paired *t*-test, $t_1$ = 6.45, *P* = 0.186, *n* = 2 mice, Holm-Sidak correction; **TC (T.S.) correlation during long-term recall relative to day 1 on B trials**: one-way RM mixed effects analysis, effect of training day, $F_{4, 5.09}$ = 14.9, **P = 0.005, *n* = 3 mice; **Day 1 vs. Day 25 B trial recall**: two-tailed paired *t*-test, $t_1$ = 4.71, *P* = 0.133, *n* = 2 mice, Holm-Sidak correction; **Day 1 vs. Day 30 B trial recall**: two-tailed paired *t*-test, $t_1$ = 23.39, *P* = 0.054, *n* = 2 mice, Holm-Sidak correction). (d) Neighboring imaging sessions remained stable as well using same analysis (**Neighboring session PV correlation during long-term recall A trials**: one-way RM mixed effects analysis, effect of training day, $F_{4, 4.35}$ = 2.69, *P* = 0.17, *n* = 3 mice; **Day 1 vs. 6 Vs. Day 20 vs. Day 25 A trial recall**: two-tailed paired *t*-test, $t_1$ = -0.43, *P* = 0.741, *n* = 2 mice, Holm-Sidak correction; **Neighboring session PV correlation during long-term recall B trials**: one-way RM mixed effects analysis, effect of training day, $F_{4, 4.01}$ = 12.91, *P = 0.015, *n* = 3 mice; **Day 1 vs. 6 Vs. Day 20 vs. Day 25 B trial recall**: two-tailed paired *t*-test, $t_1$ = -3.3, *P* = 0.339, *n* = 2 mice, Holm-Sidak correction; **Neighboring session TC (S.I.) correlation during long-term recall A trials**: one-way RM mixed effects analysis, effect of training day, $F_{4, 4.36}$ = 4.56, *P* = 0.076, *n* = 3 mice; **Day 1 vs. 6 Vs. Day 20 vs. Day 25 A trial recall**: two-tailed paired *t*-test, $t_1$ = -0.57, *P* = 0.672, *n* = 2 mice, Holm-Sidak correction; **Neighboring session TC (S.I.) correlation during long-term recall B trials**: one-way RM mixed effects analysis, effect of training day, $F_{4, 4.25}$ = 1.95, *P* = 0.259, *n* = 3 mice; **Day 1 vs. 6 Vs. Day 20 vs. Day 25 B trial recall**: two-tailed paired *t*-test, $t_1$ = -14.44, *P* = 0.086, *n* = 2 mice, Holm-Sidak correction; **Neighboring session TC (T.S.) correlation during long-term recall A trials**: one-way RM mixed effects analysis, effect of training day, $F_{4, 4.7}$ = 0.85, *P* = 0.551, *n* = 3 mice; **Day 1 vs. 6 Vs. Day 20 vs. Day 25 A trial recall**: two-tailed paired *t*-test, $t_1$ = 0.33, *P* = 0.96, *n* = 2 mice, Holm-Sidak correction; **Neighboring session TC (T.S.) correlation during long-term recall B trials**: one-way RM mixed effects analysis, effect of training day, $F_{4, 4.47}$ = 0.44, *P* = 0.775, *n* = 3 mice; **Day 1 vs. 6 Vs. Day 20 vs. Day 25 B trial recall**: two-tailed paired *t*-test, $t_1$ = -1.88, *P* = 0.526, *n* = 2 mice, Holm-Sidak correction). (e) No decorrelation in place maps was observed between A and B trial laps for matching neurons across time with either place tuning criterion (**Day one normalized A vs. B lap correlation (S.I.) scores for matching neurons**: Kruskal-Wallis test, $H_4$ = 6.9, $P$ = 0.141, $n$ = 786 neurons from 3 mice; **Day 6 recall**: two-tailed 1-sample Wilcoxon signed-rank test against 1, $W_{160}$ = 1365, $P$ = 0.249, $n$ = 161 neurons; **Day 25 recall**: two-tailed 1-sample Wilcoxon signed-rank test against 1, $W_{172}$ = -1361, $P$ = 0.302, $n$ = 173 neurons; **Day one normalized A vs. B lap correlation (T.S) scores for matching neurons**: Kruskal-Wallis test, $H_4$ = 3.68, $P$ = 0.451, $n$ = 251 neurons from 3 mice; **Day 6 recall**: two-tailed 1-sample Wilcoxon signed-rank test against 1, $W_{58}$ = 16, $P$ = 0.952, $n$ = 59 neurons; **Day 25 recall**: two-tailed 1-sample Wilcoxon signed-rank test against 1, $W_{61}$ = -319, $P$ = 0.263, $n$ = 62 neurons).

### Extended Data Fig. 1

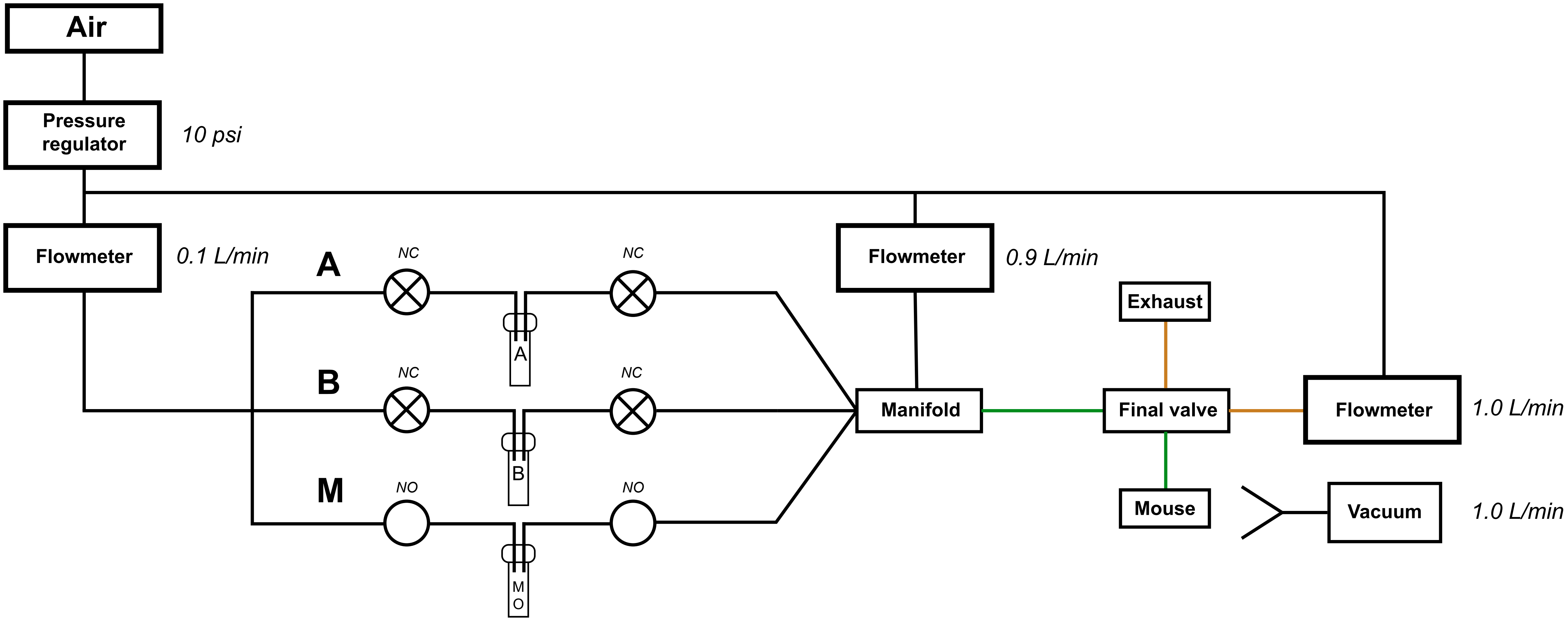

Extended Data Fig. 2

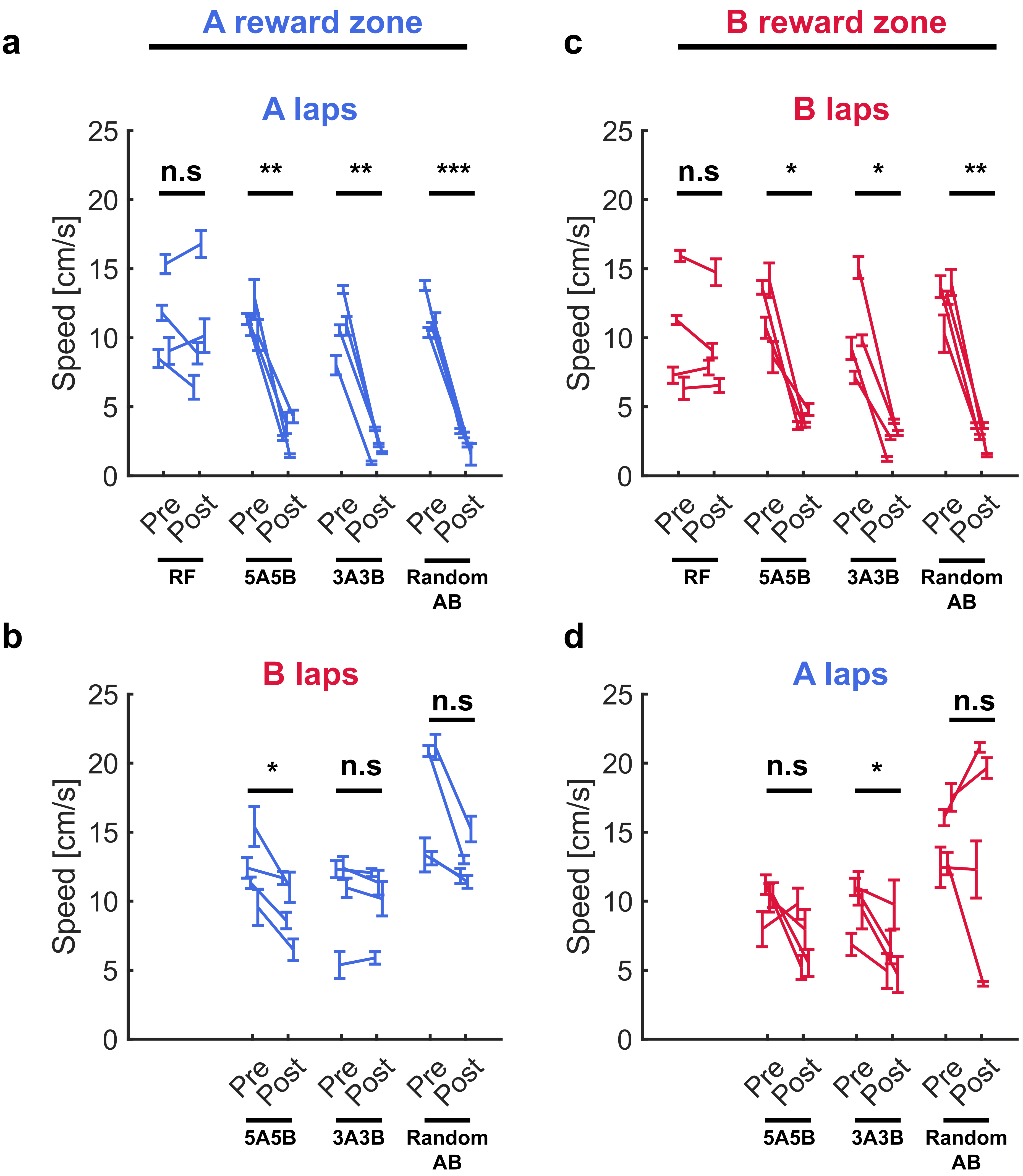

### Extended Data Fig. 3

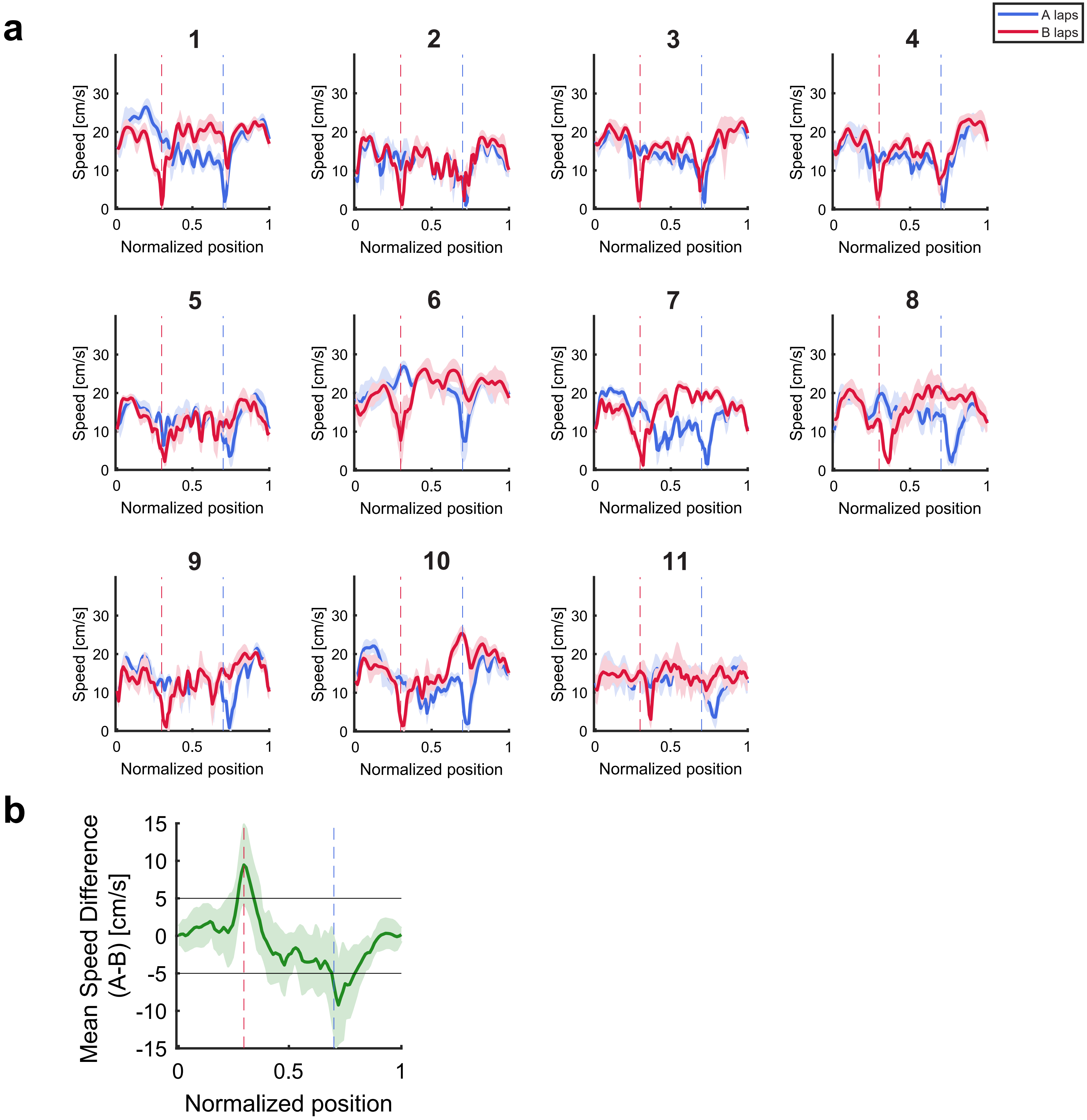

### Extended Data Fig. 4

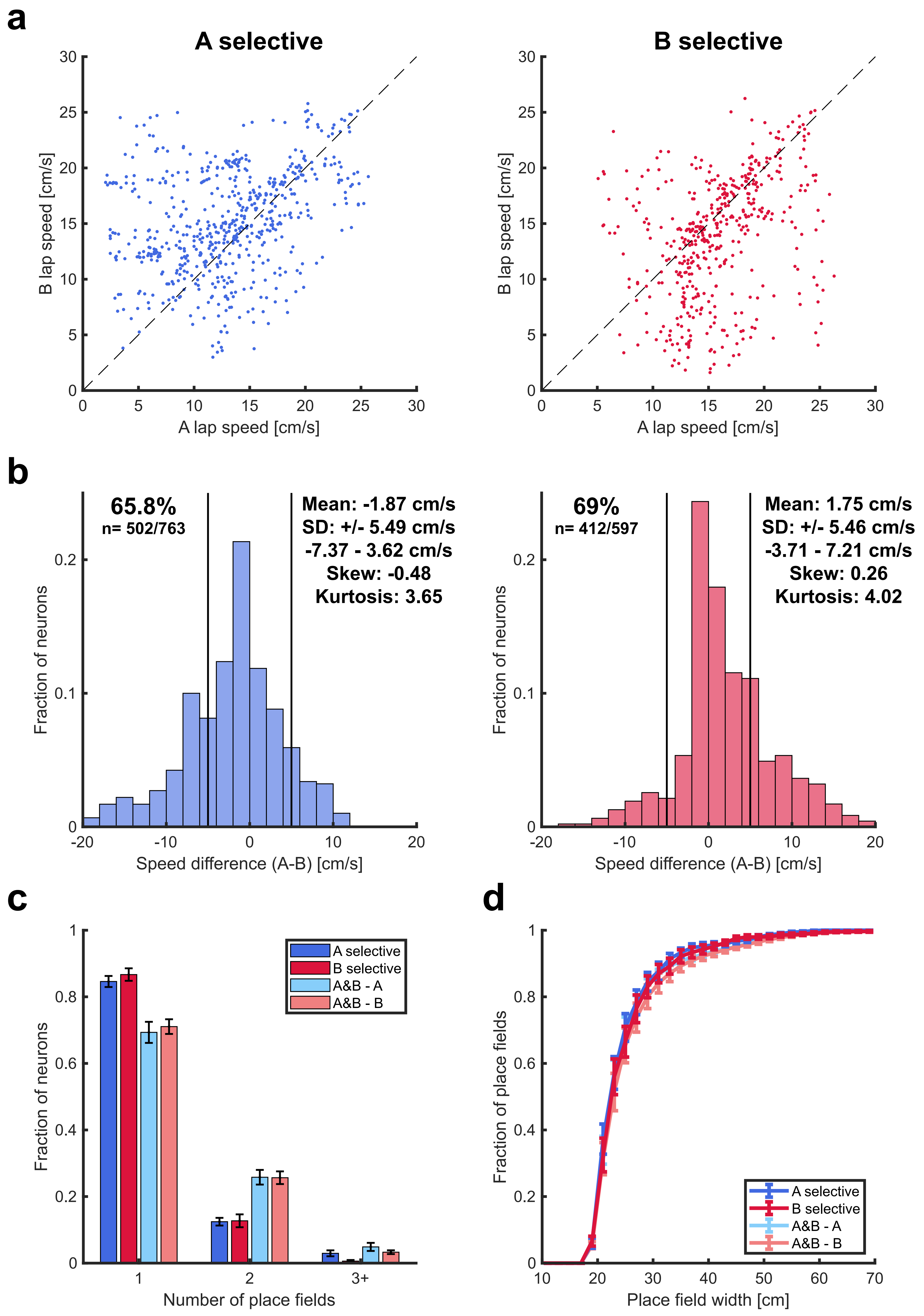

#### Extended Data Fig. 5

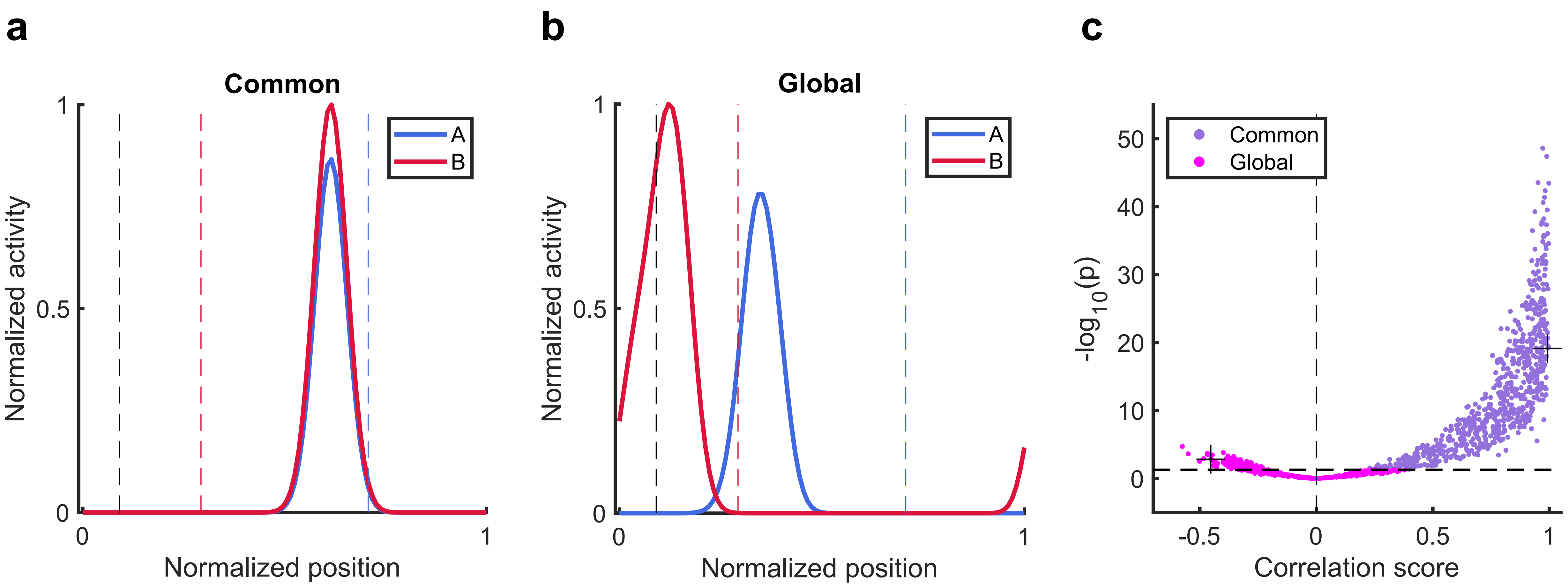

### Extended Data Fig. 6

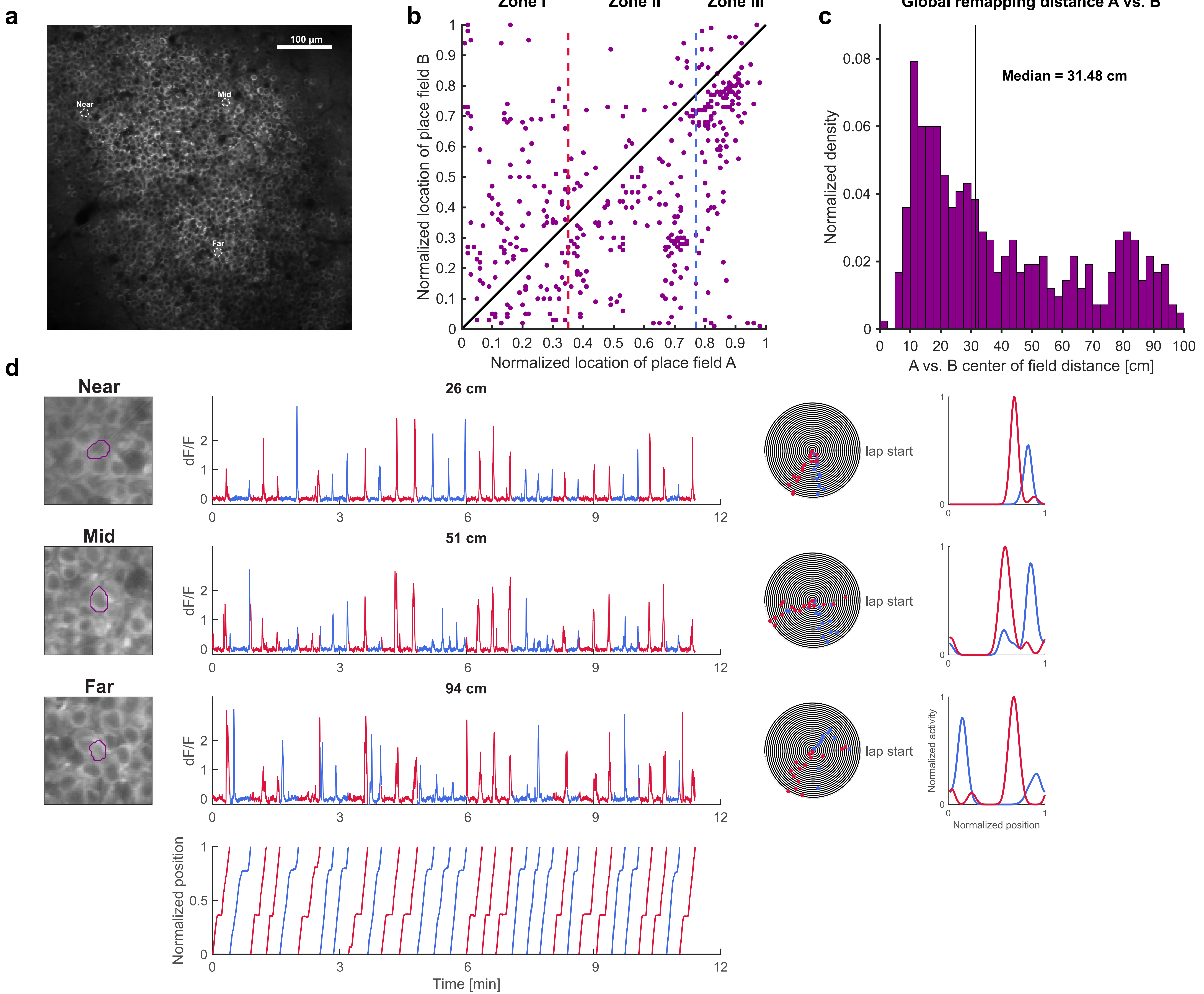

### Extended Data Fig. 7

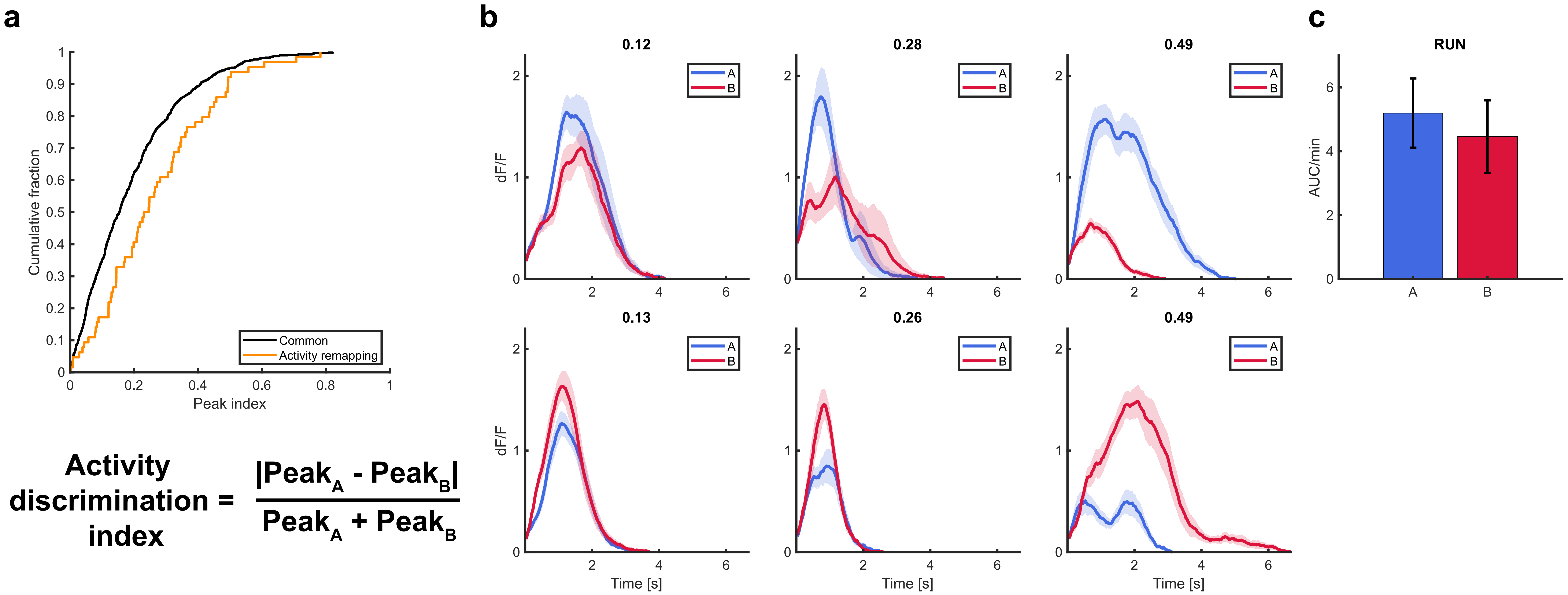

#### Extended Data Fig. 8

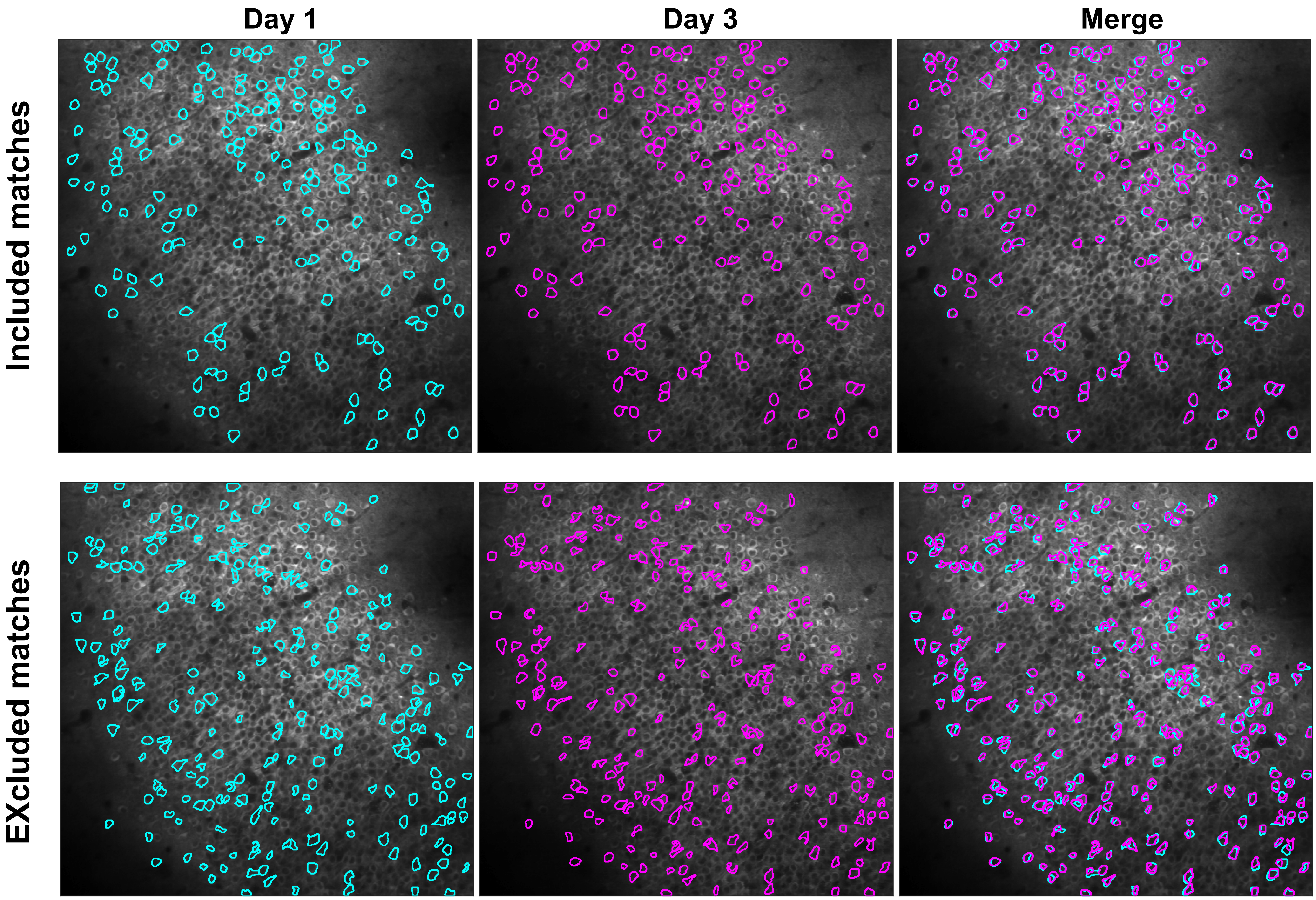

#### Extended Data Fig. 9

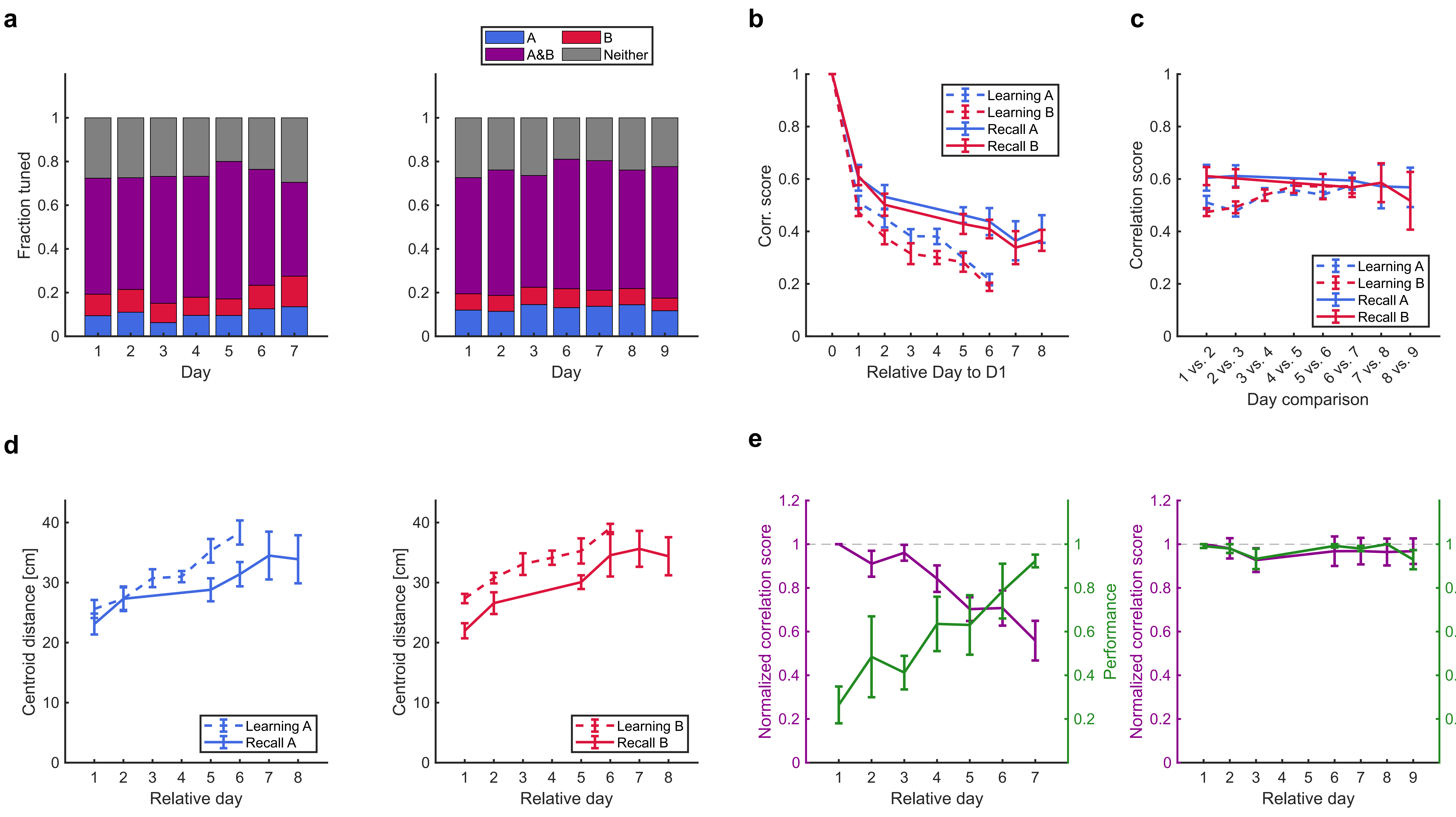

### Extended Data Fig. 10

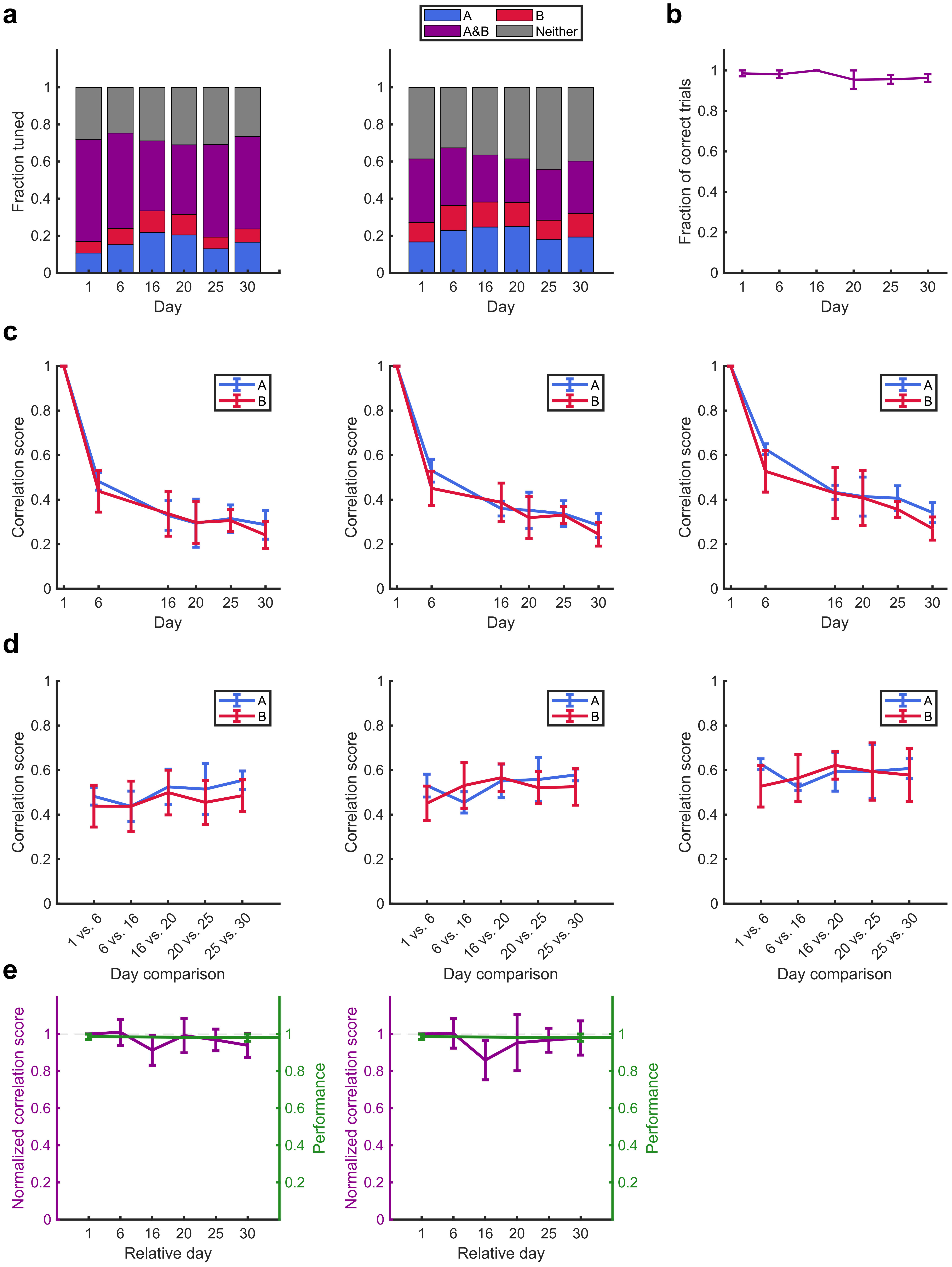
